# Working Memory Representations in Visual Cortex Mediate the Effects of Distraction

**DOI:** 10.1101/2021.02.01.429259

**Authors:** Grace E. Hallenbeck, Thomas C. Sprague, Masih Rahmati, Kartik K. Sreenivasan, Clayton E. Curtis

**Affiliations:** Department of Psychology, New York University, New York, NY 10003; Center for Neural Science, New York University, New York, NY 10003; Department of Psychological and Brain Sciences, University of California, Santa Barbara, CA, 93106; Division of Science and Mathematics, New York University Abu Dhabi, Abu Dhabi, UAE

**Keywords:** working memory, attention, fMRI, modeling, human, retinotopy, saccades

## Abstract

Although the contents of working memory (WM) can be decoded from activity in visual cortex, these representations may play a limited role if they are not robust to distraction. Here, we used model-based fMRI to estimate the impact that a distracting visual task had on WM representations in several visual field maps in visual and frontoparietal association cortex. Distraction caused the fidelity of WM representations in all maps to briefly dip when both the memorandum and distractor were jointly encoded by the population activities. Moreover, distraction induced small biases in memory errors which were predicted by biases in neural decoding in early visual cortex, but not other regions. Although distraction briefly disrupts WM representations, the widespread redundancy with which WM information is encoded may protect against catastrophic loss. In early visual cortex, nonetheless, the neural representation of information in WM and behavioral performance were intertwined, solidifying its importance in memory.

## Introduction

Higher cognition depends on working memory (WM), the ability to maintain and perform operations on internal representations of information no longer available in the environment. Early studies focused on a primary role of prefrontal cortex in supporting WM representations based on sustained activation over delay periods (Curtis and D’Esposito, 2003; Funahashi et al., 1989; Fuster and Alexander, 1971; Miller et al., 1993). However, recent studies have repeatedly demonstrated that the contents of WM can be decoded from the multivariate patterns of activity in early visual cortex (Christophel et al., 2012, 2017; Harrison and Tong, 2009; Jerde et al., 2012; Riggall and Postle, 2012; Serences et al., 2009; Sprague et al., 2014). This putative dichotomy between regions coordinating WM processes, located in a frontoparietal network, and regions storing remembered information, located within the sensory cortex, has led to the development of the *sensory recruitment model* of WM (Albers et al., 2013; Christophel et al., 2017; Curtis and D’Esposito, 2003; Emrich et al., 2013; Harrison and Tong, 2009; Postle, 2006; Rahmati et al., 2018; Riggall and Postle, 2012; Serences, 2016; Serences et al., 2009; Theeuwes et al., 2005). This model hypothesizes that feedback signals from frontoparietal cortex recruit the mechanisms used for perceptual encoding in sensory cortex for precise memory storage.

Yet, the idea that visual cortex plays a critical role in working memory continues to be met with skepticism (Leavitt et al., 2017; Xu, 2017, 2020), and major aspects of the sensory recruitment theory remain underspecified. For instance, it is unclear how neural circuitry in early visual cortex is able to simultaneously maintain WM representations while encoding incoming percepts (Xu, 2017). To address this, several recent studies have evaluated how WM representations might be maintained in the presence of behaviorally irrelevant sensory distraction. In each of these studies, participants remembered a visual stimulus over a delay while simultaneously viewing an irrelevant visual stimulus. The results are highly inconsistent. WM representations in visual cortex were eliminated (Bettencourt and Xu, 2016, Exp. 1), disrupted (Rademaker et al., 2019, Exp. 2), biased (Lorenc et al., 2018), or spared (Rademaker et al., 2019, Exp. 1; (Bettencourt and Xu, 2016, Exp. 3) when viewing different types of distracting stimuli under slightly different conditions. Because visual cortex does not always contain a representation of remembered information during distraction, its role in WM representation has been considered unimportant (Xu, 2017).

In contrast, the contents of WM remain decodable from parietal cortex activation patterns even when those in visual cortex are lost (Bettencourt and Xu, 2016; Lorenc et al., 2018; Rademaker et al., 2019; Xu, 2020), and in some cases the parietal WM representations are stored in a different format from that used to represent sensory information (Rademaker et al., 2019, Exp. 2). Based on these results, one might conclude that parietal cortex, and not visual cortex, maintains distractor-resistant memory representations. But, in addition to the inconsistency of the results in visual cortex, several outstanding issues prevent such a conclusion. For instance, behavioral performance on WM tasks can be biased by distracting stimuli, suggesting that interference between the mechanisms used for perceptual encoding and memory storage may underlie these effects (Magnussen and Greenlee, 1992; Magnussen et al., 1991; Rademaker et al., 2015; Smyth, 1996; Smyth and Scholey, 1994). In these behavioral studies, only distracting stimuli that are closely matched to remembered stimuli impact behavioral recall performance (e.g., viewing an irrelevant oriented grating while remembering an oriented grating). Additionally, maintaining information in visual WM impacts visual perception and visual motor selection in a manner consistent with low level sensory interference (Gayet et al., 2013; Hollingworth et al., 2013). However, previous studies have not been able to link behavioral evidence of distraction to distortions in encoded WM representations, as distraction either had no impact on behavior (Bettencourt and Xu, 2016; Rademaker et al., 2019, Exp. 1) or only a group-level worsening (Rademaker et al., 2019, Exp. 2) or biasing of WM performance (Lorenc et al., 2018) following distraction. If the neural representation of WM in visual cortex is disrupted by distraction while behavior is spared, then its role in WM is likely to be very limited, for instance to artificial laboratory conditions with simple blank retention intervals. On the other hand, if biases in WM performance were predicted by biases in neural WM representations in some regions, this would demonstrate a crucial role for the WM representations carried by those regions. Currently, critical evidence linking behavioral performance on individual WM trials and the neural representation of information within WM is lacking. Here, we tested the hypothesis that distractor-induced distortions in WM may stem from corresponding distortions in the most critical population-encoded WM representations in the visual and association cortices.

Participants precisely maintained position of a single WM target over a 12 s delay period, which previous studies show are encoded in retinotopically-organized visual field maps (Jerde et al., 2012; Rahmati et al., 2018; Saber et al., 2015; Sprague et al., 2014). The distractors, placed at controlled geometric distances from the WM targets, provided a feedforward stimulus drive while requiring a voluntary withdrawal of attention from the WM target so participants could make a difficult perceptual discrimination. We reasoned that such a distractor should have effects both at lower (e.g., visual cortex) and higher (e.g., association cortex) levels impacting both memory storage and control functions. These task features enabled us to measure WM representations before, during, and after distraction along with conjoint distractor representations. Consistent with prior reports, we found that during distraction, the fidelity of WM representations briefly dipped in visual, parietal, and frontal cortex. However, the WM representation quickly recovered before the end of the delay period. Critically, the offset in represented spatial location in recovered WM representations in visual cortex, but not in parietal cortex, correlated with trial-by-trial biases in WM behavior, demonstrating that WM reports may depend on the read-out of these specific WM representations. Together, these results point to a primary role for sensory regions in WM maintenance, supporting a key prediction of sensory recruitment theory.

## RESULTS

### Attention Drawn to a Distractor Impacts Working Memory Performance

Participants performed a memory-guided saccade task in which they precisely remembered the location of a target on each trial (12° eccentricity, random polar angle; Figure 1A-B). On 70% of trials, a distractor task occured in the middle of the memory delay where participants reported the direction of rotational motion of a random dot motion stimulus. To test our main hypothesis, we performed a series of analyses designed to estimate the effect of distraction on WM. First, we asked if the distractor impacted the quality of WM (Figure 1C-D). Memory-guided saccades were less precise on distractor-present compared to distractor absent trials (Figure 1D, *p*=0.039; two-tailed *t*-test). Similarly, the initiation of memory-guided saccades was slower on distractor-present trials (Figure 1E; *p*=0.011; two-tailed *t*-test). The magnitude of these distrator-induced perturbations to memory-guided saccades were similar to those induced by transcranial magnetic stimulation applied to frontal and parietal cortex (Mackey et al., 2017). Moreover, as previous work indicates that distractors that are more similar to memoranda are most disruptive (Magnussen and Greenlee, 1992; Magnussen et al., 1991; Pasternak and Greenlee, 2005; Rademaker et al., 2015; Smyth, 1996; Smyth and Scholey, 1994), we tested whether the impact of distraction depended on how close the distractor was to the WM target. These follow-up analyses found no significant evidence that WM precision (*p*=0.310; one-way repeated-measures ANOVA) or response times (*p*=0.149; one-way repeated-measures ANOVA) varied by their locations relative to the WM target on distractor-present trials (Figure S1A). Overall, we observed that a brief behaviorally-relevant distractor impacted WM performance similar to previous studies that reported unattended distractors interfered with WM performance (Lorenc et al., 2018; Rademaker et al., 2015; Rademaker et al., 2019, Exp. 2; but see Rademaker et al., 2019, Exp. 1, and Bettencourt and Xu, 2016 for reports that distraction has no effect on WM performance).

**Figure 1.**
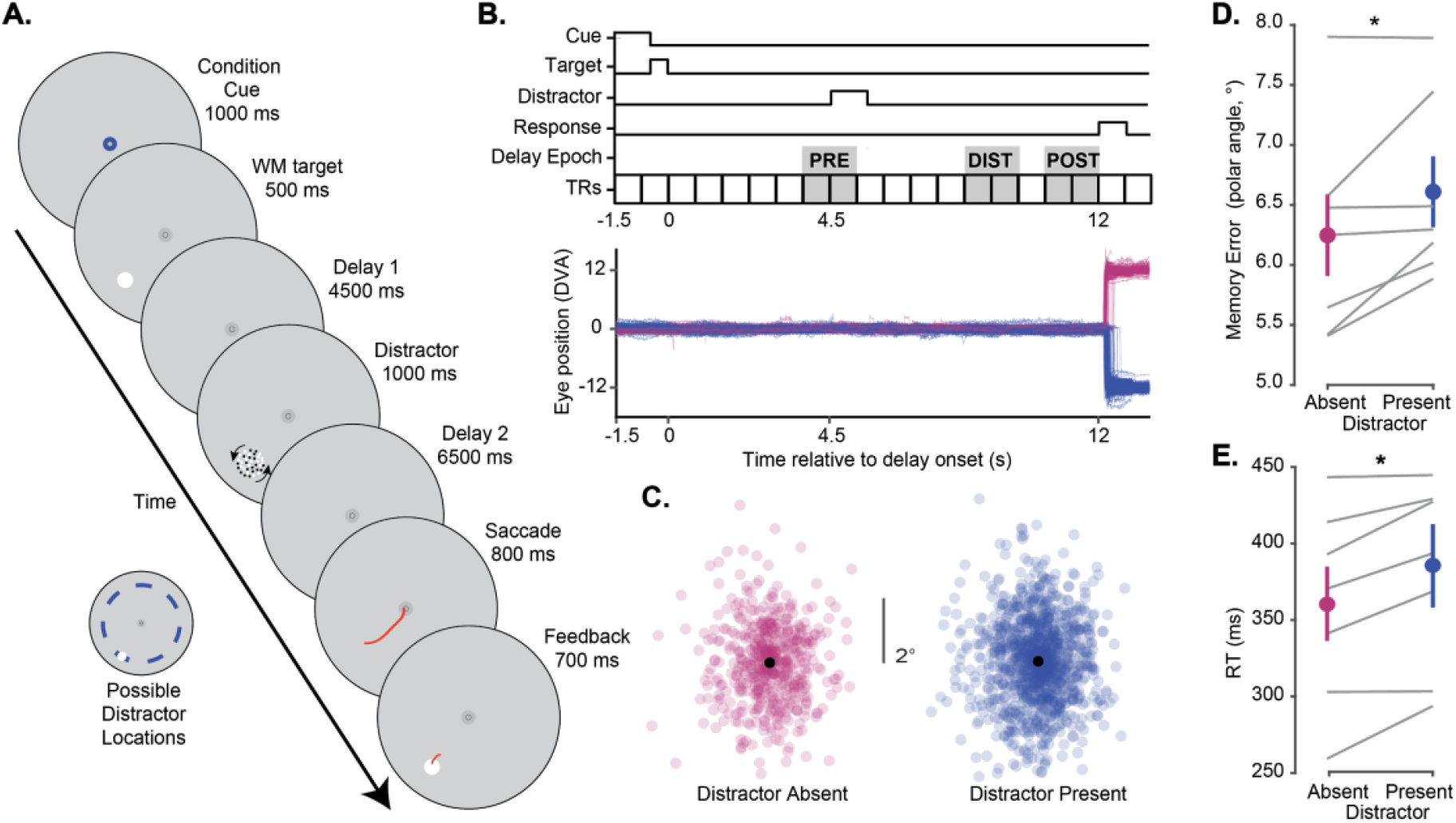
Attending a distractor stimulus impairs working memory performance. **A**. Participants (n = 7) performed a memory-guided saccade task while brain activity and gaze were recorded inside the scanner. Each trial began with a condition cue, reliably indicating whether a distractor would appear (70% of trials) or not (30%). On each trial, participants maintained the precise spatial position of a briefly-presented visual target (12° eccentricity, random polar angle) over an extended 12 s memory delay. At the end of the delay, they executed a memory-guided saccade to the remembered position. The memory target was then re-presented, and participants fixated this location before returning to central fixation. During distractor-present trials, participants discriminated whether dots presented within a 2° diameter aperture were rotating clockwise or counterclockwise with a button press. Across runs, motion coherence was varied to achieve ∼75% correct performance (mean ± SEM, 73%). The distracting stimulus could appear within one of seven position bins (24° polar angle wide) around the screen relative to the WM target, evenly presented across trials, denoted by blue intervals relative to an example WM target position (inset). **B**. Timing of task events and example gaze data. Top: Trial events (start of delay, distractor, response) were synchronized to the beginning of 750 ms imaging volumes. We defined three trial epochs for further analyses (Figures 5-7) assuming ∼4 s hemodynamic delay (PRE: volumes before distractor, DIST: volumes during distractor, POST: volumes after distractor). Bottom: Eye-trace of all trials of each condition for an example participant (p02). Eye position eccentricity is plotted as a function of time; distractor-absent trials are plotted with positive values, and distractor-present trials are plotted with negative values. Note that gaze remains at fixation during distraction keeping the retinal position of the memory target constant. **C**. Aligned final saccadic endpoints (all participants) for trials in which distractors were absent or present. All endpoints are aligned by rotating to a common spatial position (along the horizontal meridian at 12° eccentricity). **D**. Memory error (standard deviation of the polar angle of saccade endpoints) varied with distractor presence (t-test, two-tailed, p=0.039). Gray lines show individual participants; colored circles show group mean (±SEM). **E**. Response time also varied based on distractor presence (t-test, two-tailed, p=0.011). Analysis of behavioral performance across individual distractor location bins shown in Figure S1.

### Spatially-Selective BOLD Activations Persist During WM and Respond to Distraction

All fMRI analyses focused on retinotopic regions of interest identified using maps of polar angle and eccentricity computed from independent retinotopic mapping data using a compressive spatial summation voxel receptive field model (Mackey et al., 2017; Figure 3A). We analyzed data from all retinotopic ROIs previously reported in fMRI studies of visual WM and distraction (V1, V2, V3, V3AB, IPS0, IPS1, IPS2, IPS3, sPCS, LO1, hV4; (Bettencourt and Xu, 2016; Rademaker et al., 2019), and combined regions which share a foveal confluence (V1, V2 and V3; IPS0 and IPS1; IPS2 and IPS3; Mackey et al., 2017; Wandell et al., 2005, 2007). To index WM-related sustained activation within voxels tuned to remembered locations, we took advantage of the receptive field parameters and averaged BOLD responses across voxels in each visual field map whose receptive field matched that of the WM target location (RF_in_). We chose RF_in_ voxels with eccentricities between 2-15 degrees and polar angles within 15 degrees on either side of the WM target’s polar angle. For comparison, we also averaged BOLD responses in the voxels with receptive fields 165-195 degrees opposite the WM target (RF_out_) using the same range of eccentricity limits. Focusing on the distractor-absent trials, we saw two patterns across the visual field map ROIs that can be seen in Figure 2A. First, the amplitude of persistent activity during the delay period increases moving anterior in the dorsal stream ROIs from early visual cortex (V1-V3; V3AB) to parietal cortex (IPS0/1, IPS2/3) to frontal cortex (sPCS), consistent with previous reports (Emrich et al., 2013; Harrison and Tong, 2009; Jerde et al., 2012; Saber et al., 2015; Serences et al., 2009; Sprague et al., 2014). Second, the spatial selectivity of the persistent activity varied among the ROIs, which is apparent when comparing the RF_in_ and RF_out_ responses. Note how the amplitudes of delay period activity between the RF_in_ and RF_out_ conditions diminish from early visual cortex, to parietal cortex, to frontal cortex. These differences are consistent with the increasing size of receptive fields of neurons as one moves up the visual hierarchy from early visual cortex to parietal and frontal cortex (Felleman and Van Essen, 1991; Mackey et al., 2017; Wandell et al., 2007). We quantified these effects by testing whether delay-period activation (averaged over the period spanning 3.75 to 12 s after delay period onset) differed across position-sorted voxels and ROIs with a 2-way repeated measures ANOVA against a shuffled null distribution with factors of ROI and RF condition (RF_in_ vs. RF_out_). The 2-way interaction was significant (*p*=0.01), as were each of the main effects of ROI (*p*<0.001) and RF (*p*< 0.001; individual ROI statistics in Table S2). On distractor-present trials, the attended distractor evoked a phasic response in all ROIs, but this response was especially strong in parietal and frontal cortex (Figure 2B). Finally, we averaged the BOLD responses in voxels whose RFs were aligned to the distractor position, regardless of the position of the WM target, to better visualize the effect of the distractor (Figure 2C). The phasic distractor responses were robust in voxels with RFs that matched (RF_in_) compared to opposite to the distractor (RF_out_), especially in early visual cortex. Next, we used these BOLD responses to model how WM targets are encoded within the population activity of each ROI, and how distraction may disrupt such encoding across the duration of each trial.

**Figure 2.**
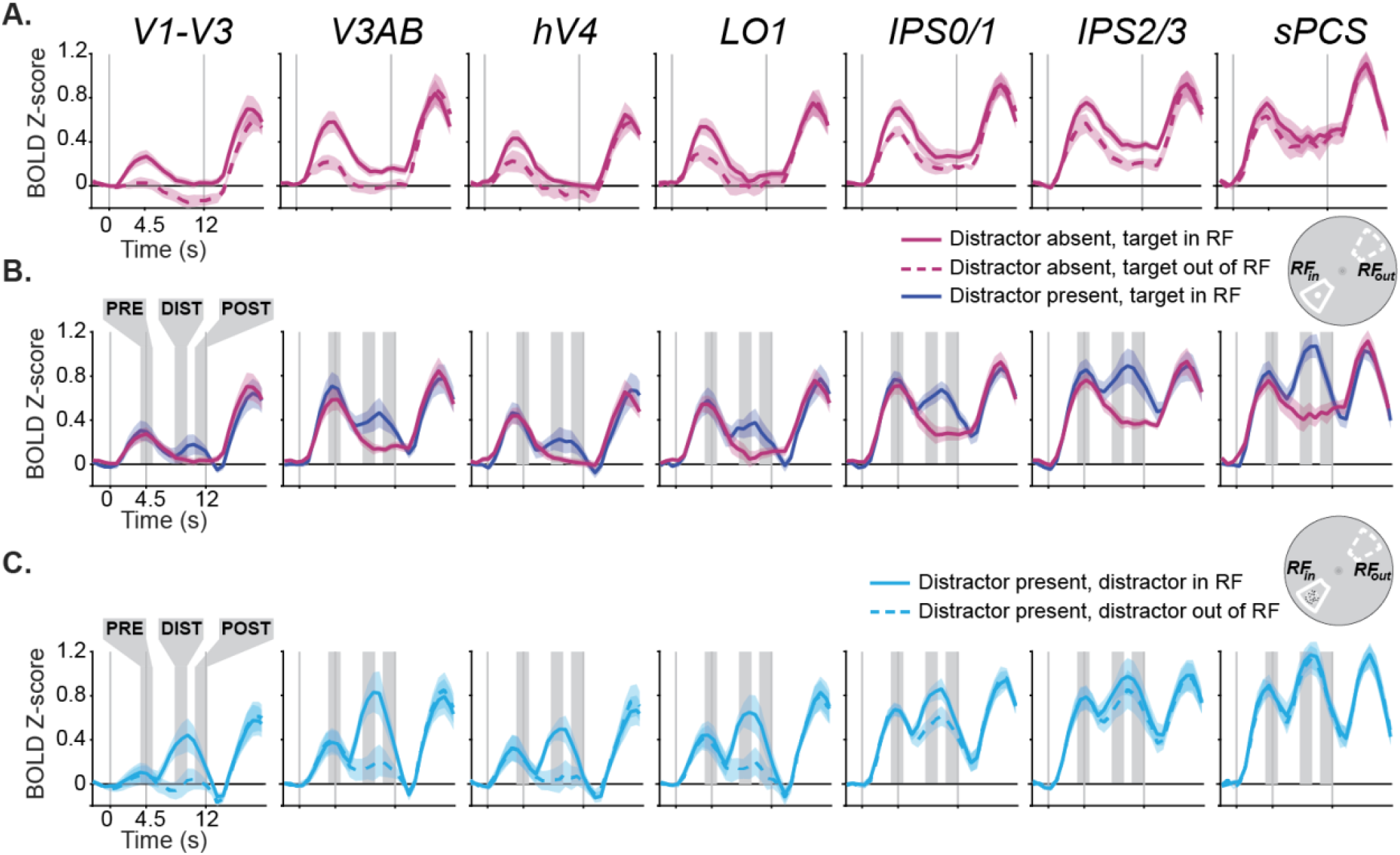
BOLD responses sorted by voxel RF position during WM delay period. **A**. During distractor-absent trials, the average (±SEM) amplitude of BOLD responses was greater in voxels whose receptive fields - estimated using nonlinear population receptive field mapping (Mackey et al., 2017, see Fig. 3A for an example hemisphere) aligned with the WM target (RF_in_) compared to when the target was 180 degrees away from voxels’ receptive fields (RF_out_). The inset to the right depicts an example of the RF_in_ and RF_out_ regions of the visual field with respect to the WM target location (see STAR Methods for more details). The amplitudes of persistent activity increased moving anterior in the dorsal stream ROIs from early visual cortex (V1-V3; V3AB) to parietal cortex (IPS0/1) to frontal cortex (IPS2/3), while the spatial selectivity (difference between RF_in_ and RF_out_) decreased. Data from ventral (hV4) and lateral retinotopic regions (LO1) is also included for completeness. Time series were baseline-corrected by removing the mean activation from -2.25 - 0 s prior to delay period onset from each time series. **B**. During distractor-present trials, we observed an additional phasic response time-locked to the distractor onset across all ROIs. **C**. To further illustrate the distractor response, we averaged the BOLD responses in voxels whose RFs were aligned to the distractor position, regardless of the position of the WM target. The phasic responses were more robust in voxels with RFs that matched (RF_in_) compared to opposite to the distractor (RF_out_). The shaded areas denote the pre-distractor, distractor, and post-distractor epochs that are the target of later analyses. Results for individual ROIs shown in Figure S2, and all p-values available in Table S1.

**Figure 3.**
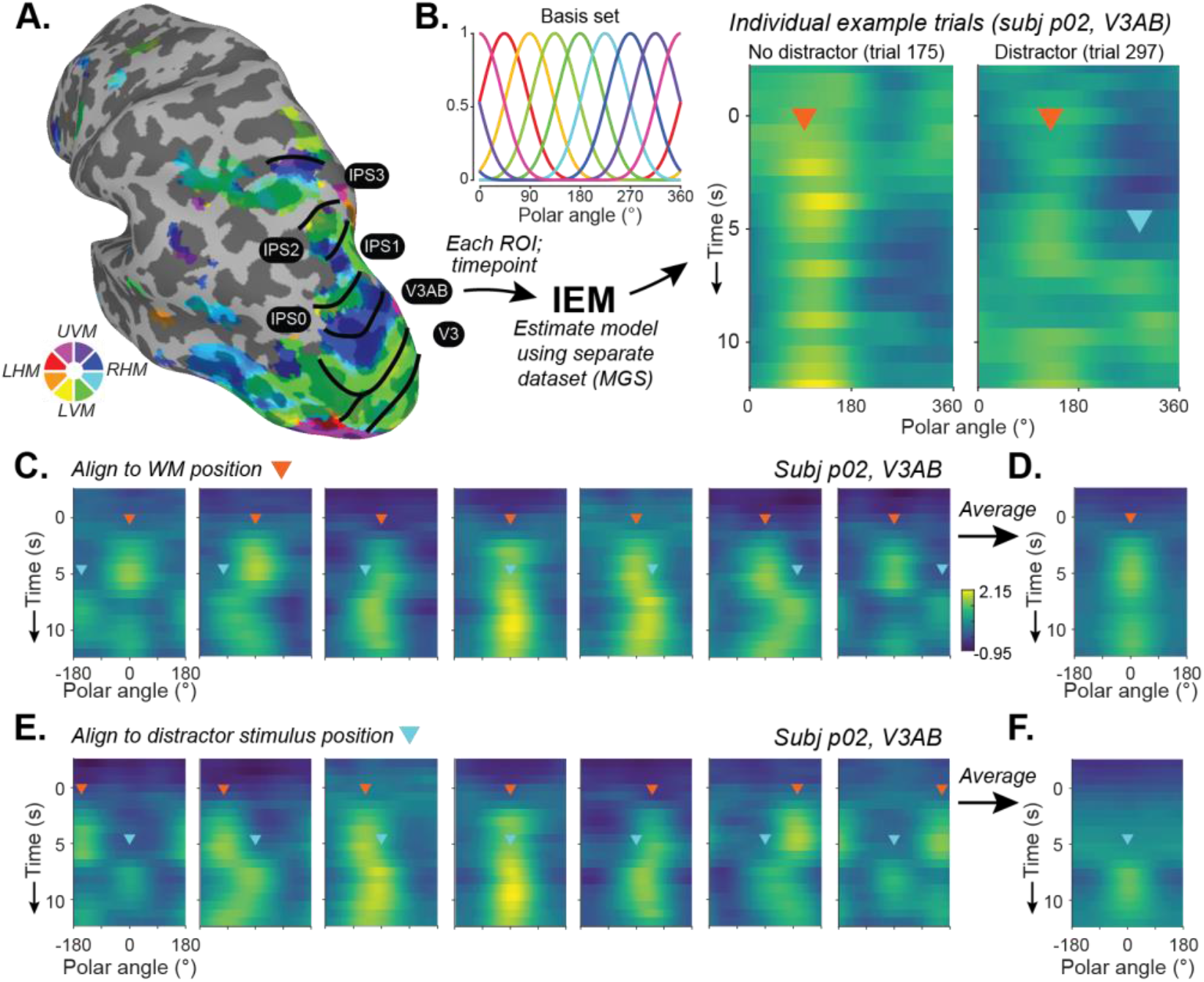
IEM-based reconstruction of WM and distractor representations. **A**. Each participant underwent retinotopic mapping to define ROIs in visual, parietal, and frontal cortex (V1-V3, V3AB, hV4, LO1, IPS0-3, sPCS). Example hemisphere and participant shown (p02, LH). Each voxel’s time course is fit with a receptive field model (Mackey et al., 2017). Color depicts preferred polar angle; thresholded at R^2^≥10%. **B**. We estimated an inverted encoding model (IEM) for polar angle for each participant and ROI using a dataset reserved for this purpose (single-item memory-guided saccade task, 3-4.5 hrs/participant). Each timepoint of the spatial distractor dataset was reconstructed using this independently-estimated model (Sprague et al., 2018a, 2019). Two example trials shown. **C**. In the example participant, reconstructions were aligned based on WM target positions (orange triangle), and separately averaged for each distractor bin position (cyan triangle at onset time). Note that both target and distractor representations can be seen in the reconstructions. **D**. WM target reconstruction averaged over all distractor location bins. Because distractors are evenly presented around the screen with respect to WM locations (Figure 1A), averaging across relative distractor positions reveals target-related spatial representations because distractor representations are ‘washed out’ during the averaging procedure. **E**. The same data as (C-D) now aligned to each trial’s distractor position (cyan triangles), and averaged separately for each relative distractor location bin (WM targets are at different locations relative to distractor; orange triangles). **F**. Distractor location reconstruction averaged over all relative WM target location bins.

### Reconstructed Population Response Encodes WM Targets and Distractors

To test our hypothesis that an attended distractor stimulus disrupts neural WM representations, we implemented an inverted encoding model (IEM) of polar angle to visualize and quantify spatial WM representations along an isoeccentric ring of target positions. We trained the encoding model using delay-period data from a single-item memory-guided saccade task collected over 2-3 fMRI sessions independent of the main experimental task. Then, using this fixed encoding model (Sprague et al., 2018a), we reconstructed 1D spatial representations from each timepoint of each trial (Figure 3A-B). Evident even in single trials, the location of the WM target emerged in the reconstructions shortly after the appearance of the stimulus and persisted throughout the entire delay period, while on distractor-present trials the position of the distractor briefly appeared and disappeared in the reconstruction (Figure 3B).

Testing our hypothesis requires visualizing WM representations independent of distractor representations. To accomplish this, we aligned reconstructions across trials based on the remembered target location, which averages over all relative distractor locations (Figure 3C-D). Similarly, to visualize the representations of the visual distractor, we aligned *the same data* based on the distractor location, which averages over all relative target locations (Figure 3E-F, as in Rademaker et al., 2019). Importantly, because distractors were presented with equal likelihood at one of seven location bins relative to the WM target (Figure 1A), we were able to independently visualize and assay WM representations and distractor representations on distractor-present trials.

Across all participants, the WM target locations were robustly encoded in the modeled population responses in all visual, parietal, and frontal ROIs during distractor-absent trials (Figure 4A). This demonstrates the robustness of our model, experimental data, and procedures, and replicates previous results (Rahmati et al., 2018; Sprague et al., 2014, 2016). Turning to the distractor-present trials, the strength of WM representations took a noticeable ‘dip’ following the distractor presentation (Figure 4B). This dip, however, was brief and the WM target representation returned shortly after the distractor disappeared, prior to the response period at the end of the trial. Additionally, by aligning the same data to the distractor location rather than the WM target location (Figure 4C), one can see strong representations of the distractor locations encoded in the population response of all ROIs.

**Figure 4.**
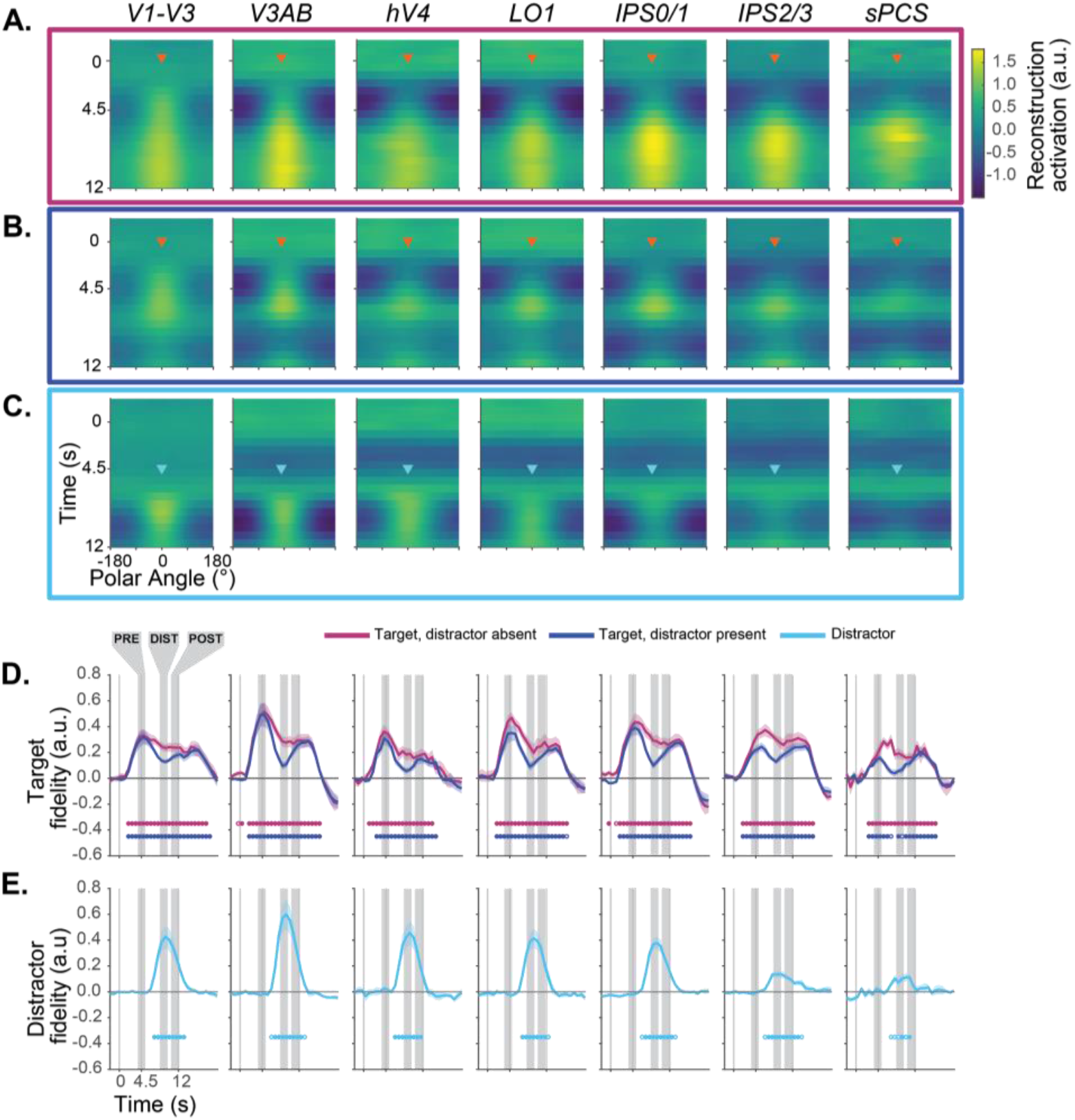
Impact of distraction on the dynamics of WM representations. Average reconstruction of WM target locations on distractor-absent (**A**) and distractor-present trials (**B**) across all participants (n=7). **C**. Reconstruction of distractor locations on distractor-present trials, where all trials were aligned to a fixed distractor location. Note that **B** and **C** include the same data, just aligned to different locations (see Fig. 3). Reconstruction strength is greatest at the aligned location in each instance, and represents the polar angle location of the WM target maintained over the entire delay period or the briefly-presented distractor. (**D & E**). Fidelity of the neural representation of WM targets (**D**) and distractors (**E**). When activation peaks in the direction of the remembered target (after alignment), fidelity is positive; when there is no consistent activation peak, fidelity is near zero. Target fidelity on distractor-absent trials is robust and statistically significant throughout the delay period in all ROIs. When the distractor is present, fidelity drops, but remains significantly above zero for all ROIs except for one 750 ms TR in sPCS. Distractor fidelity is also statistically significant in all regions, and is qualitatively most robust across extrastriate visual cortex (e.g., V3AB). Closed and open circles denote significance of p<0.05, one-sided, FDR corrected and p<0.05, one-sided, FDR uncorrected, respectively (one-sample t-test using null distribution derived from shuffled IEM; see STAR Methods). Error bars ±SEM.

### Distraction Impacts WM Representations

Thus far, qualitative examination of WM reconstructions suggests WM target representations are transiently disrupted by an attended distractor stimulus across visual, parietal, and frontal cortex. To quantify and test this hypothesis, we computed a ‘fidelity’ metric (Rademaker et al., 2019; Rahmati et al., 2020; Sprague et al., 2016), which measures the strength of the representation within the reconstruction. Reconstructions of correct WM target locations result in larger positive fidelity values, while poorly matched reconstructions produce low values. Importantly, fidelity can be computed independently for both the WM target and distractor locations because their relative locations were carefully counterbalanced and distributed (i.e., the average distractor location relative to an aligned WM target location has a fidelity of zero). On distractor-absent trials, the fidelity values in all ROIs grew significantly time-locked to the WM target presentation and remained above chance throughout the delay indicating accurate and sustained WM target encoding (Figure 4D). On distractor-present trials, we observed a phasic increase in fidelity values for the distractor location time-locked to the onset of the distractor in all ROIs (Figure 4E). Moreover, the fidelity of the WM target representation dropped in all ROIs in the time-points corresponding to the phasic distractor response (Figure 4D). Nonetheless, even during this dip in target fidelity, the values for the WM target remained significantly greater than zero indicating that the population response still contained information about the WM target location (Figure 4D-E, one-sided *t*-test at each timepoint compared against a null distribution computed with a model estimated using shuffled trial labels; FDR-corrected within each ROI).

### Recovery From the Effects of Distraction

To further characterize the temporal dynamics of how WM representations are impacted by and recover from distraction, we computed reconstructions and associated fidelity values corresponding to epochs before, during, and after distraction (Figure 5; see Figure 1B and 4D-E for epochs). First, to test for differences in WM target fidelity across ROIs, task conditions, and trial epochs, we performed a 3-way repeated-measures ANOVA against a shuffled null distribution with factors of ROI, epoch, and condition (see Methods: statistical procedures). The 3-way interaction was significant (*p*=0.041), as well as 2-way interactions of epoch × condition (*p*=0.001), epoch × ROI (*p*<0.001), and additionally, main effects of epoch, condition, and ROI (*p*<0.001; *p*=0.001; *p*<0.001). These results suggest that attending a distractor stimulus differentially impacts the WM representation across ROIs and across the delay period. Following up on the 3-way ANOVA and motivated by the dip in fidelity time-locked to the distractor onset (Figure 4D), we performed a 2-way repeated-measures ANOVA (factors of condition and epoch) within each ROI to formally test if the distractor disrupted the fidelity of the WM target representations. We observed a significant main effect of condition in all ROIs (all *p’s* <0.037), main effect of epoch in V1-3, V3AB, hV4, LO1, and IPS0/1 (all *p’s*<0.014), and interaction in V1-V3, V3AB, and IPS0/1 (all *p*’s<0.005; tests FDR-corrected across ROIs; p-values computed against a shuffled null; see Methods; Figure 5B).

**Figure 5.**
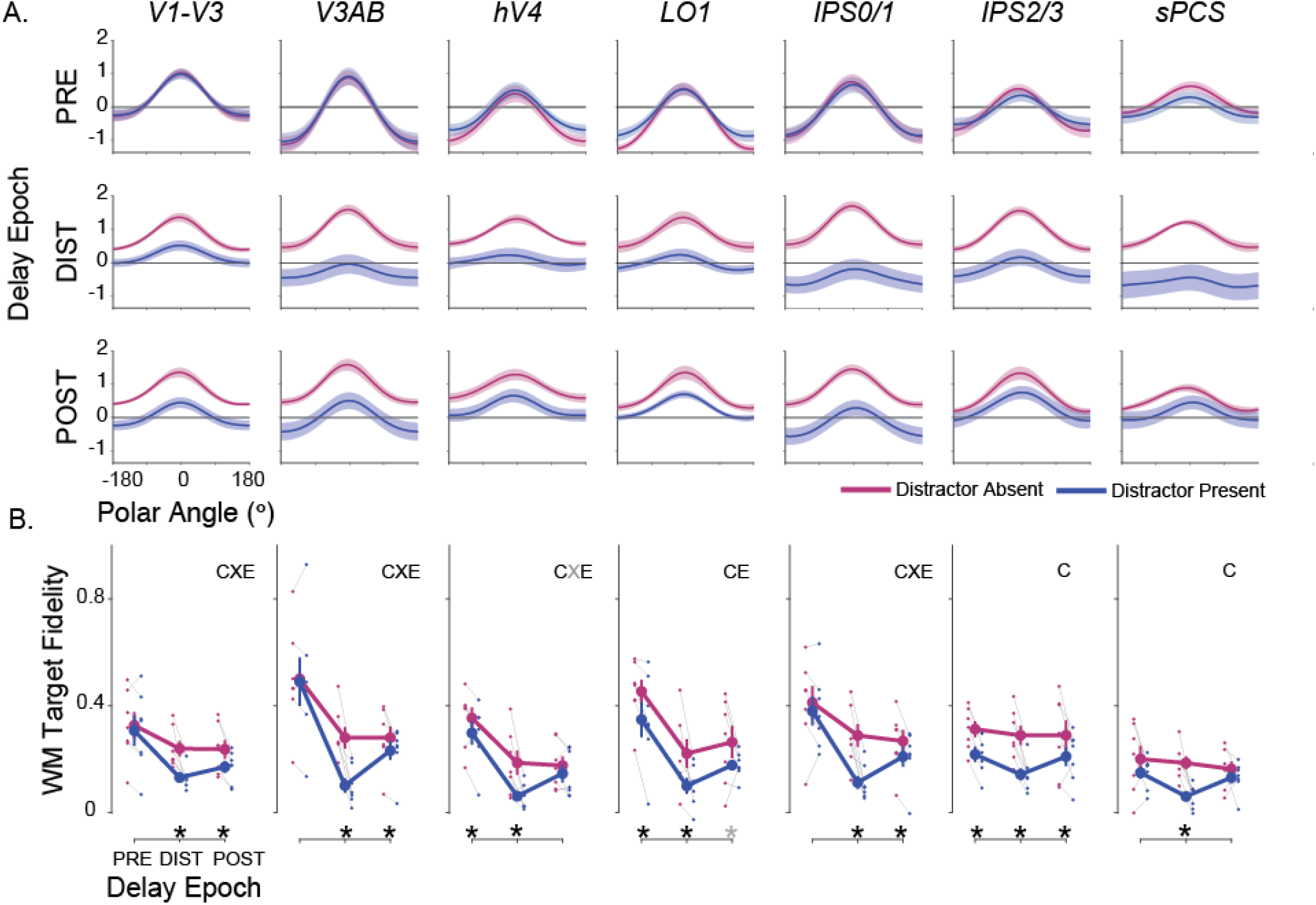
WM representations are transiently disrupted by an attended distractor. **A**. Independently trained model-based reconstructions of the WM target locations on distractor-absent (magenta) and distractor-present trials (blue) each averaged over three epochs of the memory delay. The epochs were composed of TRs before the distractor (3.75-5.25s), during the distractor (8.25-9.75s), and after the distractor (10.5-12s), accounting for the hemodynamic delay. Error bars ±SEM. Note that during the distractor epoch, the reconstructions of the WM target locations appear weaker on distractor-present compared to distractor-absent trials. In some regions, this effect of the distractor lasts into the post-distractor epoch. **B**. Average (±SEM) fidelity of reconstructed WM targets on distractor-absent (magenta) and distractor-present (blue) trials separately for the pre-distraction, distraction, and post-distraction epochs. Thin gray lines connect mean distractor-absent (small magenta dots) and distractor-present (small blue dots) fidelity for individual participants for each delay epoch. The results from 2-way ANOVAs for each ROI (epoch and condition as factors; compared against a shuffled null) are marked by symbols to denote the significant main effects of condition (C), epoch (E), and the interaction between epoch and condition (X). The significant results of paired t-tests between distractor-present and distractor-absent reconstructions per epoch, for each ROI, are marked with asterisks. In both cases, gray symbols denote p<0.05, uncorrected, and black p<0.05, FDR corrected across ROIs within test. Results for individual ROIs shown in Figure S4, and all p-values available in Table S4.

To unpack these ANOVAs, we performed follow-up *t*-tests within epochs comparing target fidelity on distractor-present and distractor-absent trials. First, WM target representations were significantly disrupted during the distractor epoch in all ROIs (Figure 5B, DIST: *p’s*<0.01; FDR-corrected across all ROIs for each epoch independently). Second, WM target fidelity remained lower following distraction compared to the same late epoch on distractor-absent trials in V1-V3, V3AB, IPS0/1, and IPS2/3 (Figure 5B, POST: *p’s*<=0.03). This suggests that the distractor degraded the quality of the WM target representations, which were not completely restored after distraction. Third, immediately following the cue that indicated whether the trial would contain a distractor, WM target fidelity was lower on distractor-present trials even prior to distractor onset in hV4, LO1, and IPS2/3 compared to distractor-absent trials (Figure 5B, PRE: *p’s*<0.027). This suggests that, when participants know a distractor will appear during a trial, they may be adopting a strategy whereby they encode WM representations in a different format during distractor-present trials than that used for distractor-absent trials (see also Bettencourt and Xu, 2016). Next, we test this possibility.

### Representational Format of WM Does Not Change During Distraction

Previous research (Bettencourt and Xu, 2016; Lorenc et al., 2018; Rademaker et al., 2019; Serences, 2016) suggests that participants may proactively insulate WM representations from the deleterious effects of sensory distraction by representing WM features in distinct formats in association cortex compared to visual cortex. Unique representational formats would thus protect at least some WM representations (those in parietal cortex) from sensory interference, while WM representations held in a static or fixed format would be susceptible to sensory interference. To test this hypothesis, we analyzed distractor-present trials using a modified model training/testing procedure. Rather than estimating a model using several independent sessions of independent mapping data, we instead trained the model using distractor-present trials only, using leave-one-run-out cross-validation. Additionally, to evaluate the possibility of a shifting/dynamic code over the delay interval (Spaak et al., 2017; Stokes et al., 2013), we repeated this procedure for each pair of timepoints during the delay period such that each time point was reconstructed with a model trained at each timepoint (King and Dehaene, 2014). If, during distraction, the dip in WM representation fidelity we observe is explained by a ‘reformatting’ or ‘recoding’ of information during distraction, this matched model training/testing procedure should show less evidence for distractor interference (Figure 6A: Stable Code). Other possible results (nonexhaustive possibilities) are qualitatively shown in Figure 6A. For example, the code may ‘morph’ following distractor presentation, resulting in a new - but incompatible - WM representation format (Figure 6A: Morphed code; Parthasarathy et al., 2017). Alternatively, if the distractor presentation transiently or permanently disrupts the WM representation, we would observe a brief (or sustained) dip in WM reconstruction fidelity, but no change in its format (Figure 6A: Stable, with transient/permanent disruption).

**Figure 6.**
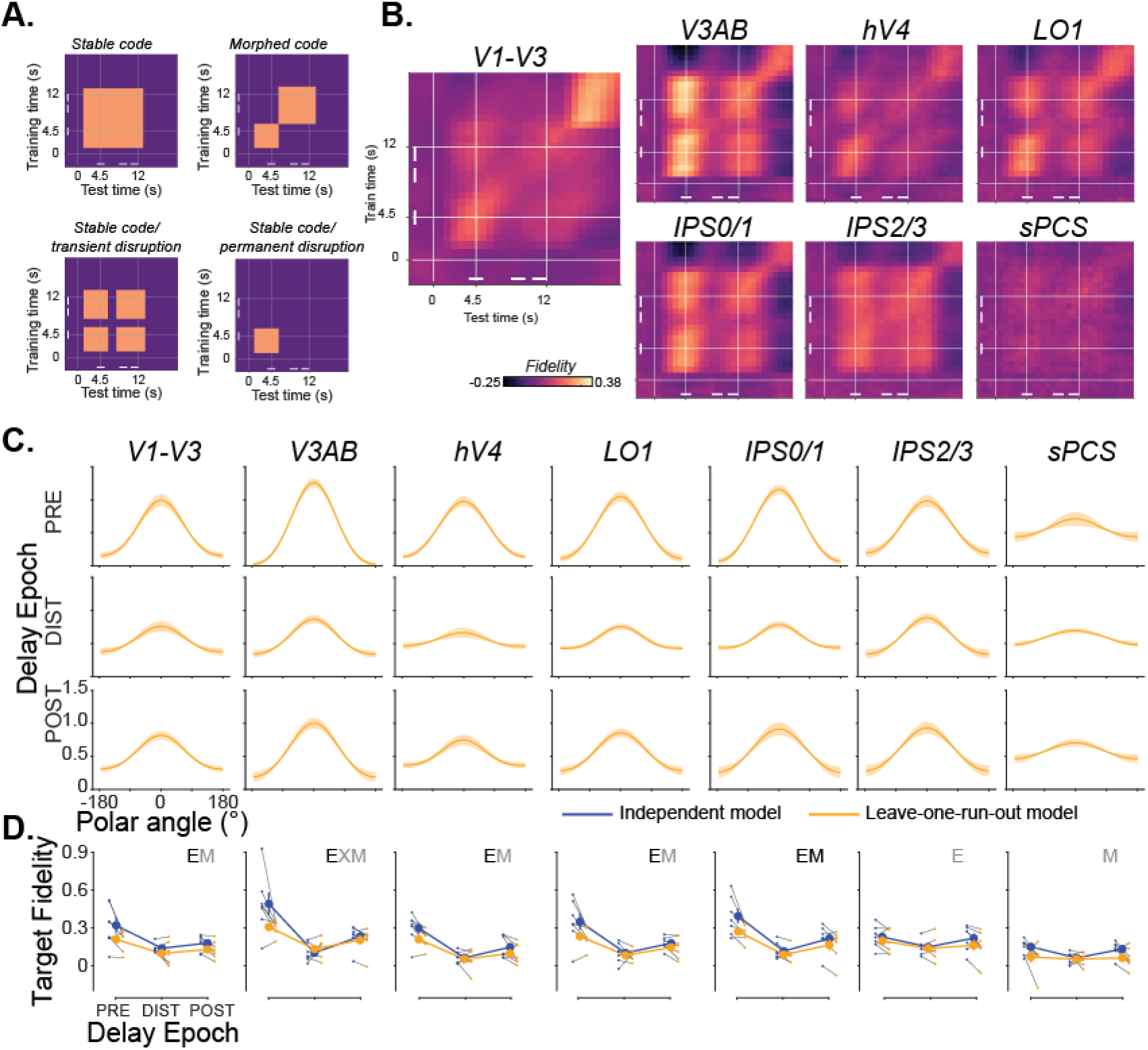
Loss of WM fidelity during distraction cannot be explained by a different coding format. **A**. To evaluate the format of WM representations throughout distractor-present trials, we conducted a temporal generalization analysis using distractor-present trials to estimate an IEM (each timepoint in turn) which was used to reconstruct held-out distractor-present trials (each timepoint in turn; leave-one-run-out cross-validation). For each combination of training and testing timepoints, we compute the WM target representation fidelity. Four cartoon examples illustrate predicted results from this analysis under various (non-exhaustive) coding schemes. **B**. Fidelity is strong across a large combination of training/testing timepoints during the delay period with no evidence of a transition to a new coding format during or after the distractor. In many ROIs (e.g., V3AB), results are consistent with a transient disruption in WM representation, but no change or morphing in representational format following distraction. White bars indicate epochs used for analyses in C-D. **C**. Model-based reconstructions from a cross-temporal generalization analysis in which training and testing was performed on corresponding epochs of the delay (i.e., train IEM with PRE timepoints, reconstruct using PRE timepoints from trials in held-out run). Rows show reconstructions from each ROI from each epoch (error bars ±SEM). Qualitatively, a substantial dip in WM reconstruction strength is apparent during the DIST epoch, as in Figure 5A. **D**. Comparison of mean fidelity during each trial epoch across model estimation procedures. Blue line shows data computed using an independent model (replotted from Figure 5B); orange line shows data computed using the leave-one-run-out cross-validation procedure. Gray lines show individual participants. We performed a 2-way repeated measures ANOVA against a shuffled null for each ROI (factors model and trial epoch). Main effects of model are indicated by M, main effects of epoch are indicated by E, and interactions are indicated by X. Significant tests are shown in black (p<0.05, FDR corrected across ROIs within test); trends are shown in gray (p<0.05, no correction). Error bars ±SEM. No ROIs show a significant interaction between model and epoch (though a trend is seen in V3AB, which is largely driven by stronger WM target representations measured using the independent model). Data from all individual ROIs available in Figure S5; p-values for all tests available in Table S6.

Using these procedures, WM target fidelity was very stable with variations in the training or testing time points having little effect on our ability to reconstruct WM representations (Figure 6B), Additionally, this approach still resulted in apparent loss of WM target information following distraction in visual (V1-V3, V3AB, hV4, LO1) and posterior parietal (IPS0/1) cortex, counter to what would be predicted if the WM representations were transiently transformed into a different format (Figure 6C-D). These results are therefore incompatible with the hypothesis that WM representations are dynamically recoded into more durable representations as a means to resist the effects of distraction, and are most aligned with the ‘Stable, with transient disruption’ model depicted in Figure 6A.

Next, we quantitatively tested whether the evolution of fidelity of WM representations over the trial differed across model estimation procedures. We computed fidelity for reconstructions generated using a leave-one-run-out cross-validation scheme where a model is trained and tested on matched timepoints (Figure 6D, orange) and compared these values to those computed using the independent model estimation procedure (same data as in Figure 5C replotted; blue). First, we performed a 3-way repeated-measures ANOVA (factors of ROI, model estimation procedure, epoch; compared against a shuffled null): there were significant main effects of ROI (*p*<0.001), model estimation procedure (*p*=0.01), and epoch (*p*<0.001), interactions between ROI × epoch (*p*<0.001) and model estimation procedure × epoch (p=0.029), and a 3-way interaction (*p*=0.018; all p-values available in Table S6). Next, we performed 2-way repeated-measures ANOVAs for each ROI (factors of model estimation procedure, epoch; compared against a shuffled null, FDR-corrected across ROIs within test). We observed a significant main effect of epoch in V1-V3, V3AB, hV4, LO1, and IPS0/1 (*p*<0.001), and a main effect of model estimation procedure in IPS0/1 (*p*=0.004). We did not observe any significant interactions between model estimation procedure and epoch, as would be predicted if WM representations are dynamically insulated from disruption via a distinct coding scheme (all *p*’s>0.079), though a trend for this interaction (*p*<0.05, no correction for multiple comparisons) was observed in V3AB. Altogether, these results do not provide evidence in support of the hypothesis that WM representations are dynamically re-coded into a novel format to protect against adverse effects of distraction.

### Distractor-Induced Offsets in Neural Representations Predict WM Errors

Our results thus far indicate that an attended distractor impacts both the quality of WM (decrease in precision and increase in RT; Figure 1D-E) and the quality of neural WM representations in visual, parietal, and frontal cortex (Figures 4-5). Next, we ask: to what extent are changes in WM performance related to distractor-induced perturbations in neural representations? We predict that memory errors (i.e., memory-guided saccade endpoints relative to the true WM target locations) will be attracted towards the location of the distractor stimulus. Moreover, we predict trial-by-trial variability in neural representations of WM targets will correlate with the direction and amplitude of memory errors. In order to test this hypothesis, we restricted our analyses to trials in which the target and distractor were near one another (within 12° polar angle), where the angle of attraction towards the distractors would align with the polar angle component of the encoding model (far distractors, instead, would predominantly align with the unmodeled eccentricity component).

We first tested whether memory errors were attracted to the nearby distractor stimulus. We quantified distractor-induced bias as the mean polar angle of memory-guided saccade endpoints flipped and rotated such that the nearby distractor was always clockwise. Indeed, WM errors were significantly biased in the direction of the attended distractor (two-tailed *t*-test compared to shuffled null, *p*=0.006; Figure 7A). Next, we tested if trial-by-trial biases in the neural representations of WM targets predicted the behavioral memory errors attracted to the distractor locations. To do so, we decoded the WM target location represented by each ROI’s WM reconstructions by taking the circular mean of the reconstruction on each trial (see Methods). Then, for each participant, we correlated each trial’s decoded WM location with the memory error on the corresponding trial (Figure 7B; all participants shown in Figure S5). Finally, we aggregated correlation values across participants and compared each ROI’s sample against 0 (one-tailed *t*-test against shuffled null). Strikingly, biases in the V1-V3 representations of the WM targets significantly predicted memory errors (*p*=0.005; corrected for multiple comparisons via FDR), but no such correlations were found in other ROIs (all *p*’s >0.104 and do not survive FDR correction; individual ROI analyses Figure 7C), and a shuffled 1-way repeated-measures ANOVA with ROI as the factor was non-significant (*p*=0.078). These results indicate that WM representations encoded in the population activity of visual cortex are not only susceptible to interference from an attention-demanding distractor task, but that the distractor distorts the neural representation causing systematic errors in memory.

**Figure 7.**
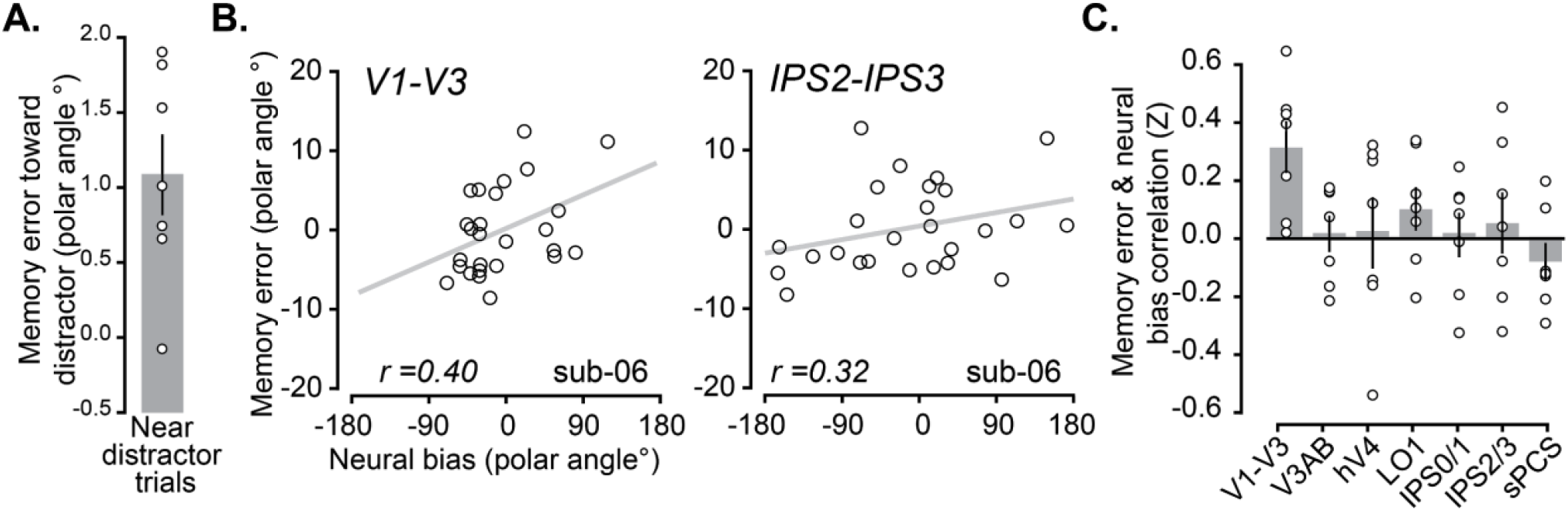
Memory errors correlate with distractor-induced biases in WM representations in visual cortex. **A**. On distractor-present trials in which the distractor was presented within 12° polar angle from the WM target, we found an attractive bias such that behavioral WM responses were drawn toward the distractor (positive values indicate errors in the same direction as distractor; two-tailed permutation t-test, p=0.006). **B**. We quantified the trial-by-trial error of each WM reconstruction based on its circular mean (see STAR Methods) during the post-distractor epoch on distractor-present trials when the distractor was presented near the WM location. To determine whether behavioral WM responses were impacted by any offsets in these neural WM representations, for each ROI and participant we correlated each trial’s decoded WM representation error with the corresponding behavioral memory error. Example scatterplots shown for one participant; trend line shows least squares linear fit (all participants and ROIs are plotted in Figure S6). **C**. Average (±SEM) neural/behavioral error correlation across participants based on decoded error from each ROI. Behavioral responses significantly correlated with errors in neural representations in V1-V3, but not other ROIs (p=0.005, FDR-corrected across ROIs; trial-level permutation test; see Methods). There was no significant main effect of ROI (p=0.08, permuted 1-way ANOVA). All p-values available in Table S8.

## DISCUSSION

We tested how distraction interferes with working memory (WM) representations to address the debated roles of early visual cortex versus association cortex. Within a widely distributed collection of visual retinotopic maps that encoded spatial WM targets, diverting attention to a salient visual stimulus caused a transient disruption, but not loss, of information about the memorandum. WM representations in parietal cortex were impacted in anticipation of distraction but showed no evidence of undergoing any transformation into a new format. Moreover, distraction caused systematic biases in memory errors, which correlated with trial-by-trial errors in the WM representations encoded in early visual cortex, but not association cortex. Based on these results, we conclude that the neural mechanisms housed in early visual cortex play a central role in WM storage, confirming key tenets of sensory recruitment theory. This conclusion that visual cortex WM representations play a critical role in WM behavior may be surprising as the functions of early visual cortex have historically been limited to visual perception. Cognitive functions like attention clearly modulate neuronal activity in visual cortex measured with fMRI (Gandhi et al., 1999; Kastner et al., 1999) or intracranial electrophysiology in humans (Martin et al., 2018) and nonhuman primates (Luck et al., 1997; Moran and Desimone, 1985; Treue and Maunsell, 1996). The sources that control attention, however, are thought to originate in frontal and parietal association cortex (Moore and Fallah, 2001). Moreover, based on electrophysiological studies, the evidence that neurons in visual cortex show persistent activity during memory delays is highly inconsistent (e.g., (van Kerkoerle et al., 2017; Mendoza-Halliday et al., 2014; Supèr et al., 2001a) recently reviewed in (Leavitt et al., 2017). Notably, WM impairments caused by PFC lesions were instrumental in establishing the link between PFC integrity and WM theory (Curtis and D’Esposito, 2004; Mackey et al., 2016), but no such evidence can be obtained in the case of visual cortex because visual impairments would confound any conclusions about memory.

The absence of evidence leaves equivocal the role of visual cortex in WM, but studies comparing the effects of distraction have yielded the most damaging evidence against the importance of WM representations in visual cortex. Neural activity in monkey inferotemporal cortex is less robust during memory delays and the selectivity of activity appears to be disrupted by intervening distractors, while PFC representations appear resistant to distraction (Miller et al., 1993, 1996). Similarly, memoranda-specific delay period activity of neurons in the monkey PFC resists the effect of distractors, unlike neurons in posterior parietal cortex (Constantinidis and Steinmetz, 1996; di Pellegrino and Wise, 1993; Qi et al., 2010; Suzuki and Gottlieb, 2013). The results from these distractor studies of single neurons are widely cited to support the claim that the PFC, rather than posterior cortical areas, is critical for WM storage. In contrast to these results, here we report that WM representations in frontal, parietal, and early visual cortex all survive distraction. The distractor does cause dips in the fidelity of the memory representations, but they are quickly restored to pre-distractor levels throughout the cortex. Indeed, the contents of WM may be protected, not by some specialized control process unique to PFC, but by the widely distributed nature of the WM representations.

More generally and in contrast to the electrophysiological studies in monkey cortex, a decade of model-based fMRI studies have demonstrated that the patterns of activity in visual cortex, including V1, can be used to decode and reconstruct the contents of WM (Albers et al., 2013; Christophel et al., 2012; Ester et al., 2009, 2013; Harrison and Tong, 2009; Rahmati et al., 2018; Serences et al., 2009; Sprague et al., 2014) recently reviewed in (Christophel et al., 2017). Moreover, memory-specific activity persists in V1 neurons (van Kerkoerle et al., 2017; Supèr et al., 2001b) and V1 voxels (Saber et al., 2015) whose receptive fields match the visual field location of WM targets (Fig. 2). Although these results together provide positive evidence in support for the sensory recruitment theory of WM (Curtis and D’Esposito, 2003; Postle, 2006; Theeuwes et al., 2005), several aspects of the theory remain controversial. One common criticism (Bettencourt and Xu, 2016; Xu, 2017) points out that, if visual cortex is maintaining WM information, how can it simultaneously process retinal input? Specifically, it remains unclear if the neural circuitry in early visual cortex is able to simultaneously maintain WM representations while encoding incoming percepts, like distracting stimuli. Also, if distraction disrupts WM representations encoded in early visual cortex without impacting the quality of WM, then visual cortex’s role in WM might be limited, for instance, to artificial laboratory experiments with blank memory delays.

Much of the recent debate regarding the impact of distraction has specifically involved the relative roles of parietal cortex and visual cortex in WM. Consistent with the previous fMRI studies of distraction (Bettencourt and Xu, 2016; Lorenc et al., 2018; Rademaker et al., 2019), visually-engaging or, in our case, attention-demanding distractors have a more reliable adverse impact on behavior and a disruptive effect on WM representations encoded in early visual cortex. Our results differ from those previously reported in several important ways. Previous studies drew conclusions based in part on the absence of significant WM decoding in visual cortex (Bettencourt and Xu, 2016; Lorenc et al., 2018; Rademaker et al., 2019). Here, we generate positive evidence that provides strong support for a critical role for visual cortex in WM. First, we find that visual cortex maintains joint representations of both sustained WM targets and transient attended distractors. These representations were impacted by, but were ultimately robust to, distraction as they were fully restored before the response period (Fig. 3, 4). Second, and most importantly, trial-by-trial distortions in these restored WM representations in visual cortex were positively correlated with errors in memory reports (Fig. 7). The positive relationship between neural representations of WM measured with BOLD fMRI and the quality of WM performance measured later in time is difficult to explain away. In contrast, the studies that have only shown disrupted WM representations during distraction rely on an absence of evidence, which may not be evidence of an absence of WM representation. Below we discuss the impact of these new findings in the context of the theoretical mechanisms by which early visual cortex and association cortex might support WM.

Previous fMRI studies suggest that WM representations in parietal cortex are resistant to distraction and therefore support the resilience of WM (Bettencourt and Xu, 2016; Lorenc et al., 2018; Rademaker et al., 2019). Presumably, parietal cortex stores WM representations in a format distinct from the sensory code used for perception, otherwise the visual input during distraction would interfere with the WM representation. Indeed, Rademaker et al. (2019) showed that parietal cortex encodes the remembered orientation, but that the representation can be in a distinct format from that of an attended oriented grating. Our results are inconsistent with this explanation. First, we find that distraction does impact WM representations in parietal cortex, if only for a brief period of time (Figs. 4-5). Second, decoding errors in the parietal representations do not predict behavior, like they do in visual cortex, indicating they may not be critically linked to the quality of WM (Fig. 7). Third, based on different training/testing procedures over time, we find no evidence that WM target representations are anything but stable codes of the visually presented WM targets (Fig. 6). Perhaps the type of memorized stimulus matters. Stimulus features that can be condensed or compressed could very well undergo recoding to minimize the demands on memory. For instance, it is unlikely that memory for dot motion is a replay of hundreds of dots moving over time, but instead a single vector representing the compressed bit of task relevant information. Therefore, the absence of motion-related activity during a memory delay in area MT is unsurprising (Mendoza-Halliday et al., 2014; Zaksas and Pasternak, 2006). A spatial location, however, may not be compressible and therefore does not necessitate any transformations.

Even if WM representations in parietal cortex are more distractor-resistant, it remains unclear why. The WM code might be less sensory in nature making it in turn less similar to incoming percepts, or control processes might actively protect their contents (Lorenc et al., 2021). In our data, we find that WM representations in parietal cortex were affected by the mere anticipation of distraction. Following the cue indicating the trial would contain distraction, the fidelity of the WM representation was lower compared to trials with no distraction. As discussed above, we ruled out the possibility that the WM representations dynamically morphed over time (Fig. 6). We hypothesize that storage of the target position and covert attention needed to rapidly detect the distractor share a common mechanism akin to the control of spatial attention. The drop in WM fidelity we observed in parietal cortex may stem from interference with the control of attention, rather than the storage of WM (Awh and Jonides, 2001; Awh et al., 1998; van Moorselaar et al., 2018; Yu et al., 2020). Regardless, WM representations in parietal cortex had no bearing on the changes in WM behavior induced by the distractor. A strong test of the importance of a neural WM representation lies in its ability to predict WM behavior. In this study, we measured neural and behavioral biases on each trial and found a strong positive correlation between the two in visual cortex alone. These results at the trial-level provide positive evidence for a critical role of visual cortex in WM, and extend previous demonstrations linking the quality of WM decoding to individual differences in WM performance (Albers et al., 2013; Christophel et al., 2018; Emrich et al., 2013; Ester et al., 2013) and differences averaged over conditions (Emrich et al., 2013; Sprague et al., 2016).

There are a number of properties that may constrain the mechanisms by which visual cortex supports WM storage. Theoretically, microcircuits that support WM through memory-specific persistent activity are supported by excitatory recurrent connections (Compte et al., 2000; Wang, 1999). The slow kinetics of NMDA receptor mediated currents in PFC support persistent activity (Wang et al., 2013), while theoretical models suggest that the faster decay rates, like those in V1 (Wang et al., 2008), would limit persistent activity (Wang, 2001). In general, intrinsic neural dynamics slow as one moves from visual, to parietal, to frontal cortex (Murray et al., 2014). Interestingly, this trend can be seen in our average BOLD time courses, where delay period activity increased systematically along the dorsal visual stream, consistent with our previous reports (Jerde et al., 2012; Saber et al., 2015). Beyond the temporal domain, relative differences in anatomical properties also suggest association cortex may have advantages over visual cortex in its capacity for WM storage. The density of NMDA receptors are less expressed in V1 than PFC (Wang et al., 2008). Pyramidal neurons in association cortex, compared to visual cortex, have larger and more complex dendritic branching with a greater number of spines (Oga et al., 2017), and have more extensive horizontal collaterals in Layers II and III (Kritzer and Goldman-Rakic, 1995). Together, these anatomical features may better equip association cortex with an increased capacity to integrate inputs, including the excitatory connections theorized to form positive feedback loops to sustain WM representations (Goldman-Rakic, 1995).

Then what role does early visual cortex play in the maintenance of WM representations? First, consider that the differences between visual cortex and association cortex in temporal dynamics and anatomy are all relative. In fact, when the recurrent network theory of WM was first proposed (Goldman-Rakic, 1995), several of the hypothesized features of PFC circuitry were unknown and were simply extrapolated from V1 (e.g., recurrence supported by horizontal connectivity between similarly tuned neurons; Gilbert and Wiesel, 1979). In theory, the same type of recurrent network could sustain WM representations in V1 as long as the rate of excitatory feedback inputs are greater than the rate of decay. Perhaps a critical source of the feedback is not local, but originates from frontal or parietal cortex. Such a mechanism is central to the sensory recruitment theory of WM, where top-down attention signals are proposed to target and boost sensory neurons to prevent memory decay (Curtis and D’Esposito, 2003). Second, the factors that presumably make visual cortex less than ideally suited for WM storage do not preclude it from being a necessary node in a larger WM network. The great precision of our visual WM likely depends on interactions between control mechanisms stemming from the association cortices and the precise encoding mechanisms in early visual cortex, and not separate systems specialized for perception and memory. Beyond WM, such concepts are supported by a growing appreciation of the critical role of visual cortex in reinstating visual percepts retrieved from episodic memory (Breedlove et al., 2020; Favila et al., 2019; Johnson and Rugg, 2007; Polyn et al., 2005; St-Yves and Naselaris, 2018) and imagery recalled from semantic memory (Pearson et al., 2015; Slotnick et al., 2005; Thirion et al., 2006).

With these ideas in mind, we designed our distractor task to not only inject noise into the population-based representation through bottom-up visual stimulation but to also interfere with the top-down signals that might be necessary to sustain WM representations in visual cortex. We found that nearby distractors had an attractive pull on memory, biasing memory errors towards the distractor, similar to previous studies (Magnussen and Greenlee, 1992; Magnussen et al., 1991; Rademaker et al., 2015; Smyth, 1996; Smyth and Scholey, 1994). Critically, when the distractor was near the WM target, fluctuations in trial-by-trial neural decoding errors in early visual cortex, but not association cortex, predicted WM errors (Fig. 7B-C). Previous studies have only reported that WM decoding errors in visual cortex predict whether distractors were clockwise or counterclockwise relative to the memoranda (Lorenc et al., 2018) and that individual differences in WM performance can be predicted by average decoding accuracy of delay period activity in visual cortex (Albers et al., 2013; Christophel et al., 2018; Emrich et al., 2013; Ester et al., 2013; Sprague et al., 2014). Our results are consistent with bump attractor models of WM that assume WM representations are self-sustained by the collective response of populations of neurons whose tuning varies along a stimulus dimension (Compte et al., 2000; Standage and Paré, 2018; Wang, 2001). Most relevant, these models predict that small random drifts in the bumps of activity cause the seemingly random inaccuracies in memory. Evidence for this hypothesis exists, as clockwise or counterclockwise biases in population estimates of delay activity in macaque PFC neurons predict small angular errors in memory-guided saccades (Wimmer et al., 2014). Using model-based fMRI in humans, we also find direct evidence to support this hypothesis but in early visual cortex, where angular decoding errors in V1-V3 predicted memory-guided saccade errors, but those in parietal or frontal cortex did not. This coupling strongly suggests that the overt report of one’s memory depends on the read-out of the population’s encoded representation in visual cortex. Ultimately, the question of whether early visual cortex is essential for visual WM, wherever one draws that line, is less relevant than trying to understand the mechanisms by which visual cortex contributes to WM.

## Acknowledgements

We thank New York University’s Center for Brain Imaging for technical support. This research was supported by the National Eye Institute (R01 EY-016407 and R01 EY-027925 to C.E. Curtis, F32 EY-028438 to T.C. Sprague, a Sloan Research Fellowship to T. C. Sprague, and by NIH T32EY007136-27 to New York University & G.E.Hallenbeck).

## SUPPLEMENTARY FIGURES

**Figure S1.**
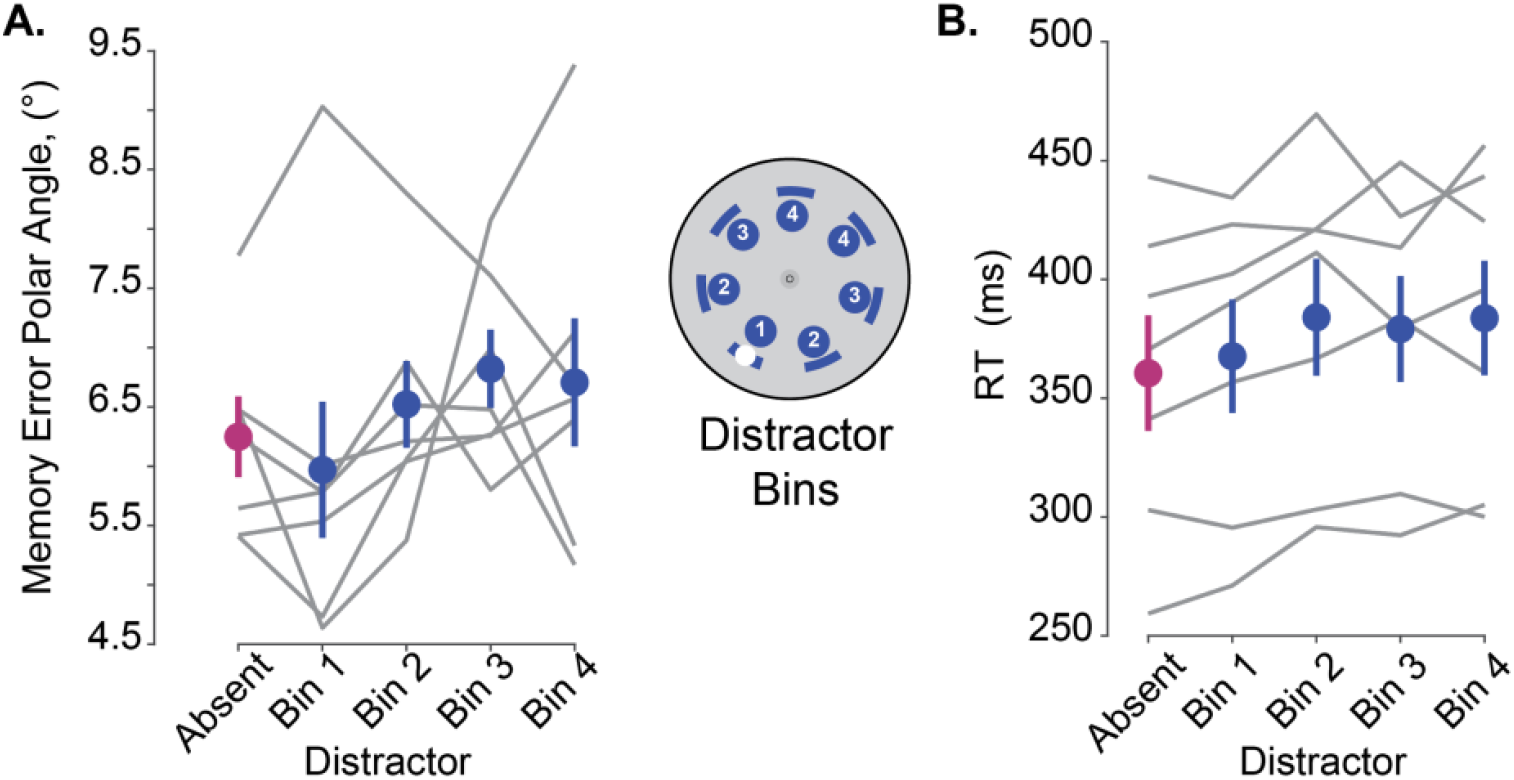
Behavioral performance does not vary across distractor offsets. **A**. Memory error on distractor absent trials (magenta) and memory error per each pseudo-randomized distractor offset bin (blue; each bin 24° wide): bin 1, 0° mean offset; bin 2, 51.42°; bin 3, 102.9°, bin 4, 154.3°, shown at inset. Gray lines depict individual participant performance across distractor-absent (30% of trials) and each distractor present condition (each offset comprised 20% of trials, with the exception of bin 1, which was 10%). To determine if the distractor-present offsets differed from one another significantly, we performed a one-way RM ANOVA on only the distractor-present conditions, and found no significant effect of memory error (*F*(3,6)=0.84, *p*=0.49). **B**. Saccadic reaction time, measured from the onset of the initial ballistic saccade from the start of the response cue period, was greater in each distractor condition (blue) as compared to distractor-absent trials. To determine whether reaction time with respect to each distractor offsets differed significantly from one another, we performed a one-way RM ANOVA. RT did not vary across distractor bins (*F*(3,6) = 2.01,*p*=0.15).

**Figure S2.**
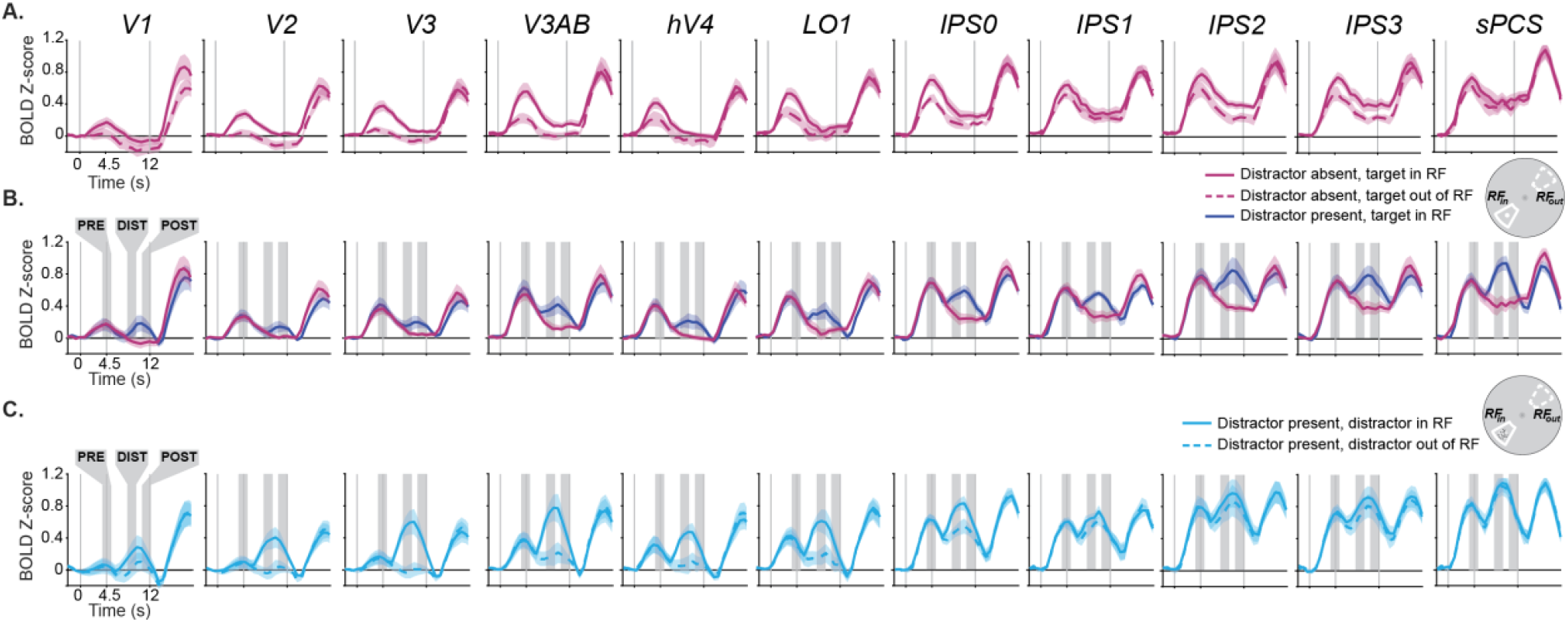
BOLD responses sorted by voxel RF position during WM delay period. **A**. During distractor-absent trials, the average (±SEM) amplitude of BOLD responses was greater in voxels whose receptive fields aligned with the WM target (RFin) compared to when the target was 180 degrees away from voxels’ receptive fields (RFout). The inset to the right depicts an example of the RFin and RFout in respect to the WM target (see STAR Methods for more details). The amplitudes of persistent activity increased moving anterior in the dorsal stream ROIs from early visual cortex to parietal cortex to frontal cortex, while the spatial selectivity decreased. **B**. During distractor-present trials, we observed an additional phasic response time-locked to the distractor onset across all ROIs. **C**. To further illustrate the distractor response, we averaged the BOLD responses in voxels whose RFs were aligned to the distractor position, regardless of the position of the WM target. The phasic responses were more robust in voxels with RFs that matched (RFin) compared to opposite to the distractor (RFout). The shaded areas denote the pre-distractor, distractor, and post-distractor epochs that are the target of later analyses.

**Figure S3.**
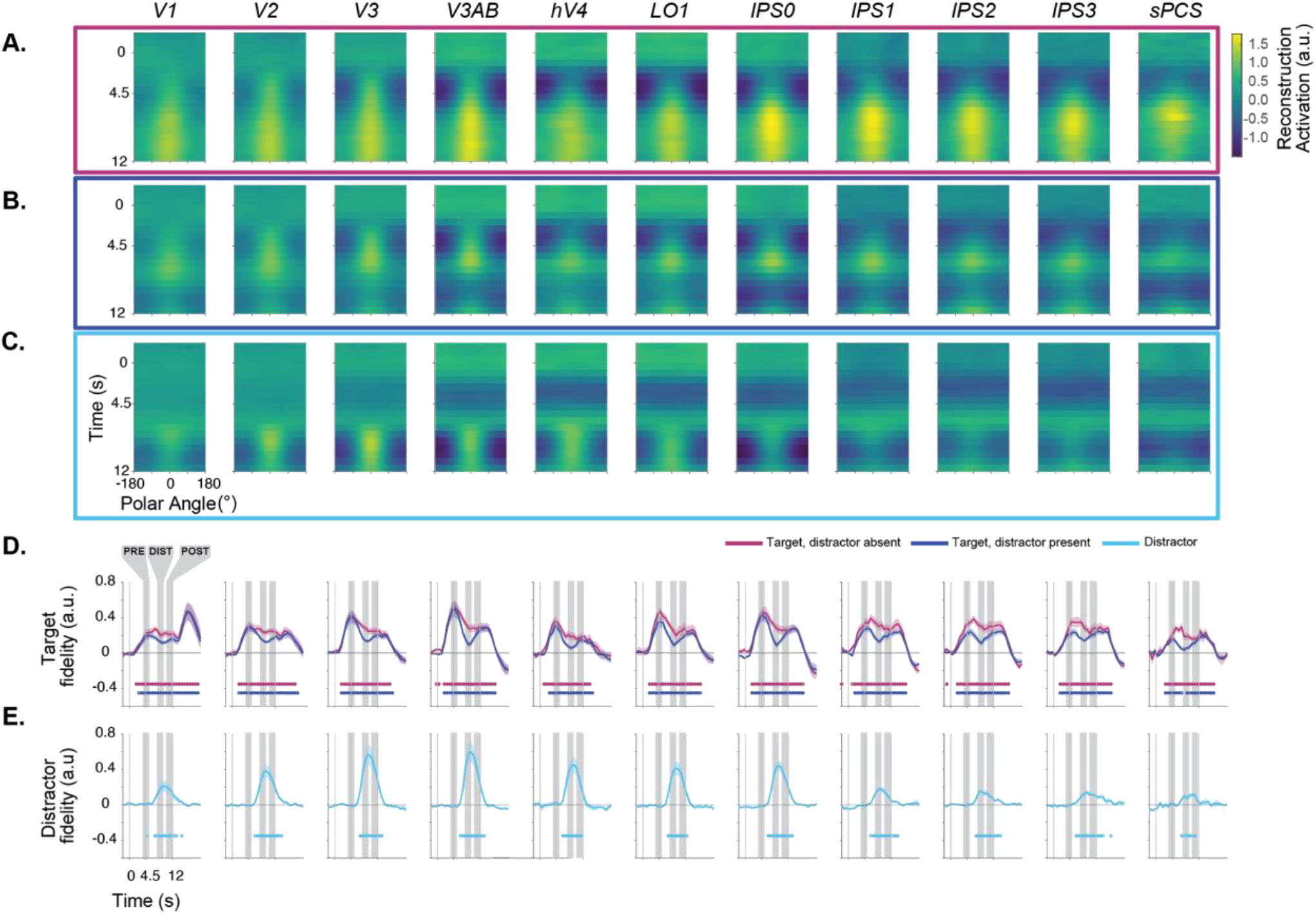
Impact of distraction on the dynamics of WM representations in individual ROIs. Data presented as in Figure 3, but for individual ROIs. Average reconstruction of WM target positions on distractor-absent trials (**A**) and distractor-present trials (**B**) across all participants (n=7). **C**. Reconstruction of distractor position on distractor-present trials, where all trials were aligned to a fixed distractor position. Note that **B** and **C** are reconstructions of the same data, just aligned to different positions. Reconstruction strength is greatest at the aligned position in each instance, and represents the polar angle position of the WM target maintained over the entire delay period or the briefly-presented distractor. (**D & E**). Fidelity of the neural representation of WM targets (**D**) and distractors (**E**). When activation peaks in the direction of the remembered target (after alignment), fidelity is positive; when there is no consistent activation peak, fidelity is near zero. Target fidelity on distractor-absent trials is robust and statistically significant throughout the delay period in all ROIs. When the distractor is present, fidelity drops, but remains significantly above zero for all ROIs except sPCS. Distractor fidelity is also statistically significant in all regions, and is qualitatively most robust across extrastriate visual cortex (e.g., V3AB). Closed and open circles denote significance of *p*<0.05, one-sided, FDR corrected and *p*<0.05, one-sided, FDR uncorrected, respectively (one-sample T-test using null distribution derived from shuffled IEM; see Methods). Error bars ±SEM.

**Figure S4.**
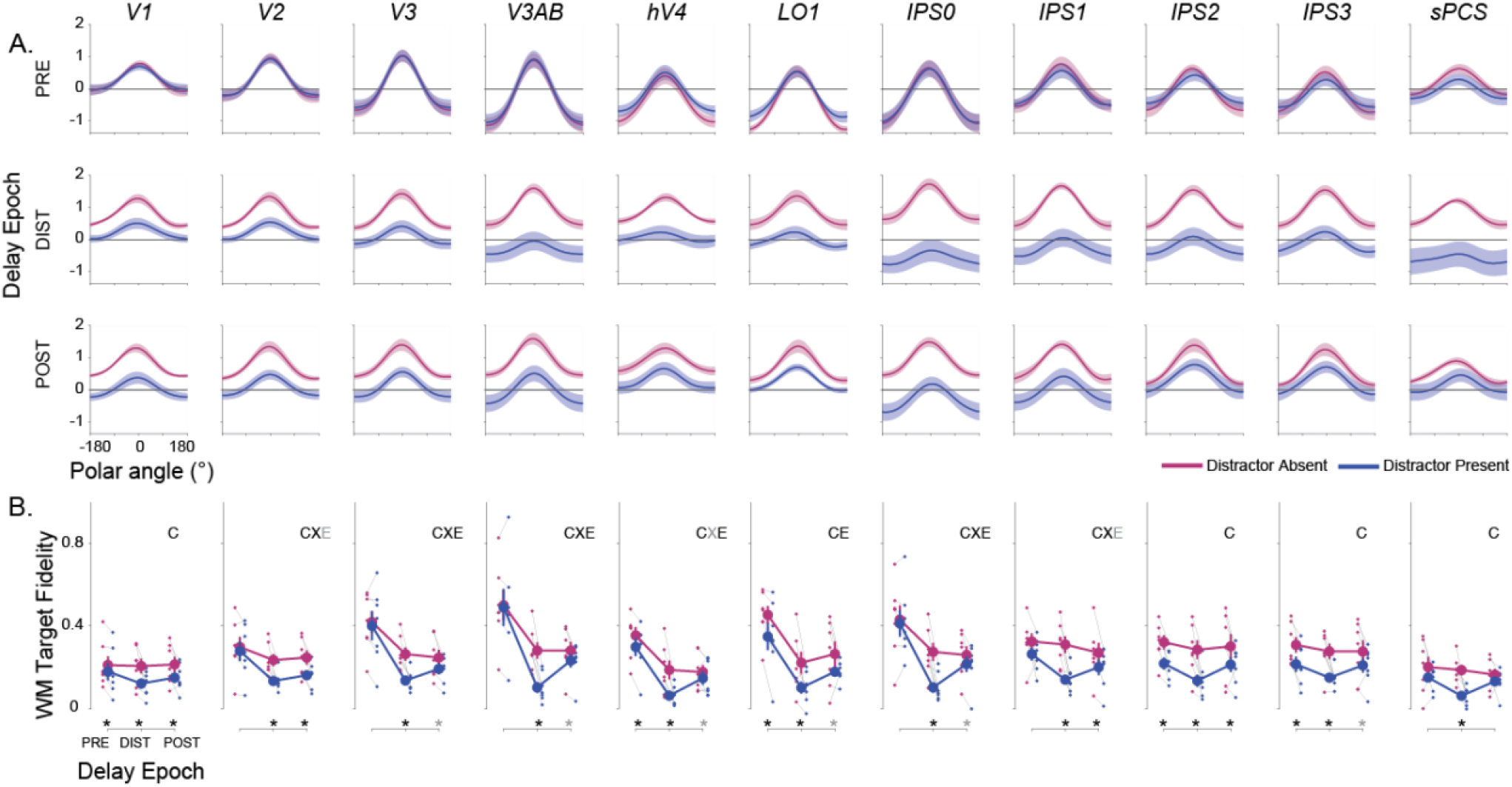
WM representations are transiently disrupted by an attended distractor stimulus in individual ROIs. Data presented as in Figure 5, but for individual ROIs. **A**. Independently trained model-based reconstructions of the WM target locations on distractor-absent (magenta) and distractor-present trials (blue) estimated separately for three epochs of the memory delay. The epochs were composed of TRs before the distractor (3.75-5.25s), during the distractor (8.25-9.75s), and after the distractor (10.5-12s). Error bars SEM. Note that during the distractor epoch, the reconstructions of the WM target locations appear weaker on distractor-present compared to distractor-absent trials. In some regions, this effect of the distractor lasts into the post-distractor epoch. **B**. Average (+/-SEM) fidelity of reconstructed WM targets on distractor-absent (magenta) and distractor-present (blue) trials separately for the pre-distraction, distraction, and post-distraction epochs. Thin gray lines connect mean distractor-absent (small magenta dots) and distractor-present (small blue dots) for individual participants for each delay epoch. The results from 2-way ANOVAs for each ROI (epoch and condition as factors; compared against a shuffled null) are marked by symbols to denote the significant main effects of condition (C), epoch (E), and the interaction between epoch and condition (X). The significant results of paired *t*-tests between distractor-present and distractor-absent reconstructions per epoch, for each ROI, are marked with asterisks. In both cases, gray symbols denote *p*<0.05, uncorrected, and black *p*<0.05, FDR corrected across ROIs. All *p*-values available in Table S3.

**Figure S5.**
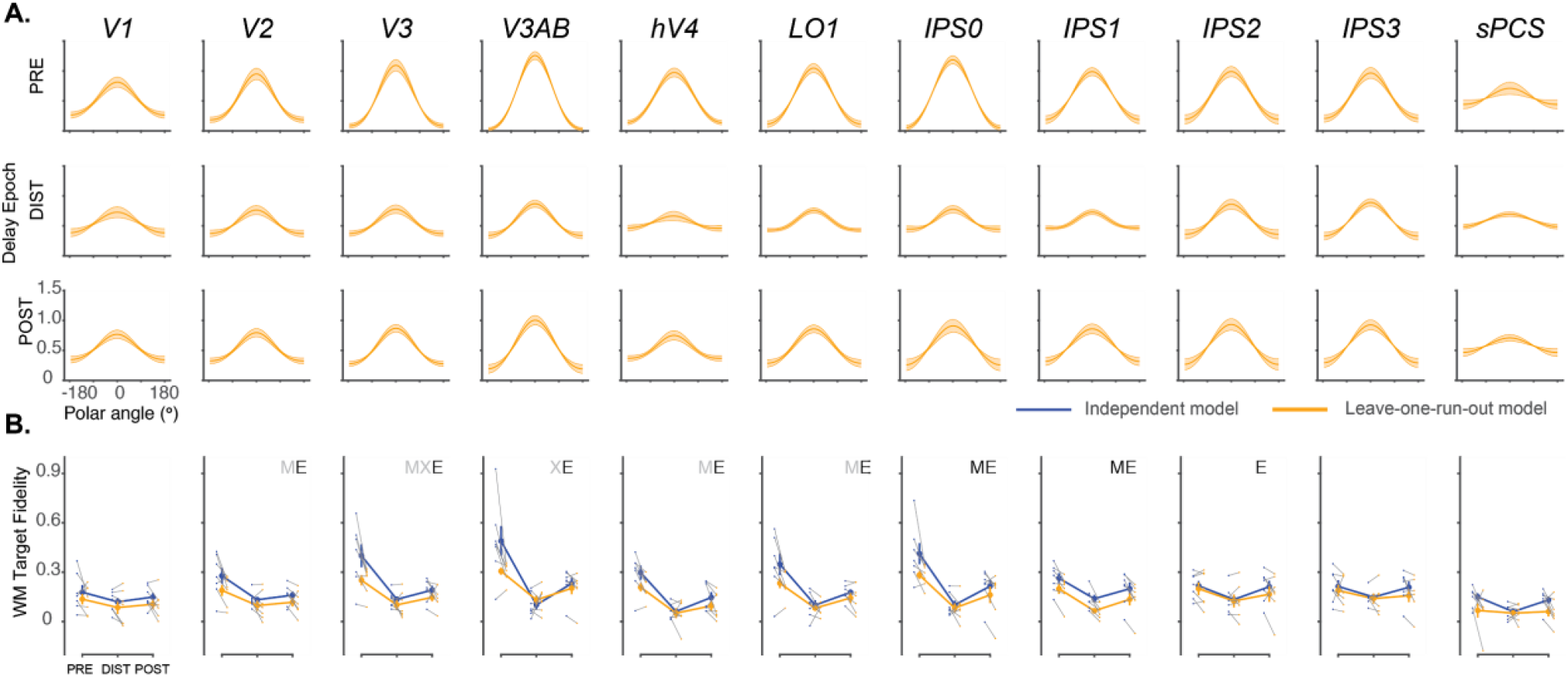
Loss of WM fidelity during distraction cannot be explained by a different coding format in individual ROIs. Data presented as in Figure 6, but for individual ROIs. **A**. Model-based reconstructions from a cross-temporal generalization analysis in which training and testing was performed on corresponding epochs of the delay (i.e., train IEM with PRE timepoints, reconstruct using PRE timepoints from trials in held-out run). Rows show reconstructions from each ROI from each epoch (error bars ±SEM). Qualitatively, a substantial dip in WM reconstruction strength is apparent during the DIST epoch, as in SFigure 6A. **B**. Comparison of mean fidelity during each trial epoch across model estimation procedures. Blue line shows data computed using an independent model (replotted from SFigure 6B); orange line shows data computed using the leave-one-run-out cross-validation procedure. Gray lines show individual participants. We performed a 2-way repeated measures ANOVA against a shuffled null for each ROI (factors model and trial epoch). Main effects of model are indicated by M, main effects of epoch are indicated by E, and interactions are indicated by X. Significant tests are shown in black (p< 0.05, FDR corrected across ROIs within test); trends are shown in gray (p<0.05, no correction). Error bars ±SEM. IPS0 & IPS0 show a significant main effect of ‘model’, though the independently trained model out-performs the leave-one-run-out model. No ROIs show a significant interaction between model and epoch (though a trend is seen in V3 & V3AB, which is largely driven by stronger WM target representations measured using the independent model). *p*-values for all tests available in Table S6.

**Figure S6.**
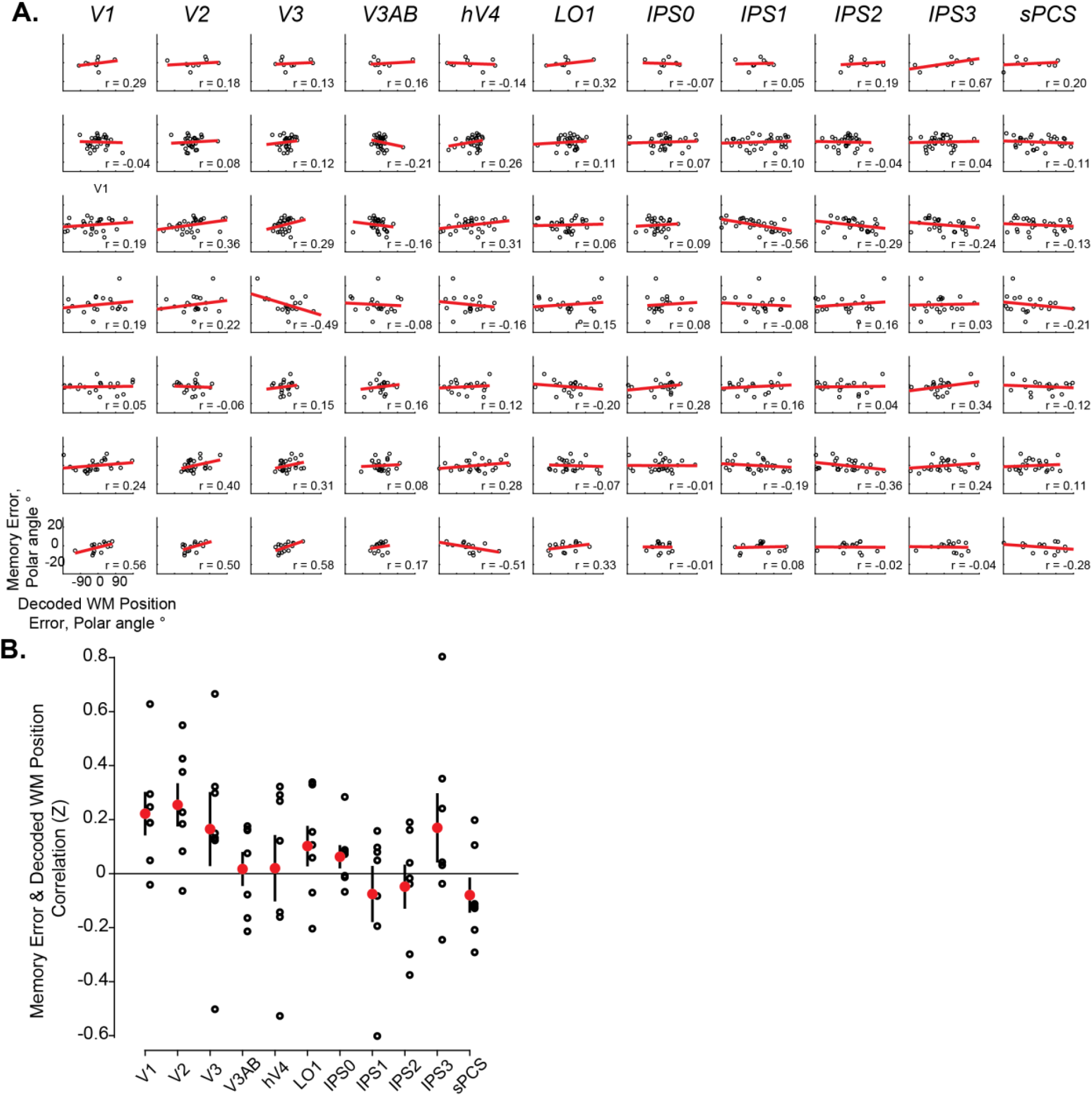
Comparison between behavioral errors and neural errors for single ROIs. **A**. Correlation between decoded WM representation error and behavioral WM response for each participant and individual ROI separately for distractor-present trials with distractor presented within 12° polar angle of WM target location. **B**. We aggregated the trial-by-trial correlation of each participants’ behavioral error with their corresponding neural error across participants (Fisher Z transformed) and compared these values against 0 (1-way *t*-test against a shuffled null). At the individual ROI level, no significant correlations were found after correction for multiple comparisons (FDR), but trends were observed in V1 and V2 (uncorrected, *p*< 0.05). A 1-way repeated-measures ANOVA did not identify a significant main effect of ROI on neural/behavioral error correlations (*p*=0.12; comparison against shuffled null). All p-values available in Table S7.

## SUPPLEMENTARY TABLES

**Table S1.** Non-parametric *p*-values characterizing differences in mean delay period amplitude and RF conditions across grouped ROIs, Figure 2. **A**. To test for differences in amplitude across ROIs and reliable differences in RF conditions across dorsal ROIs, a 2 -way shuffled ANOVA was performed on mean delay-period amplitude from distractor-absent trial with ROI and RF conditions as main effects. **B**. A follow-up 1-way shuffled ANOVA was performed within each ROI with RF condition (in vs. out) as factors.

| ROI | RF | ROI x RF |
| --- | --- | --- |
| <0.001 | 0.001 | 0.035 |

**B.**
| ROI | RF |
| --- | --- |
| V1-V3 | 0.005 |
| V3AB | 0.019 |
| hV4 | 0.009 |
| LO1 | 0.012 |
| IPS0/1 | 0.016 |
| IPS2/3 | 0.016 |
| sPCS | 0.008 |
| FDR Thresh | 0.019 |

**Table S2.**
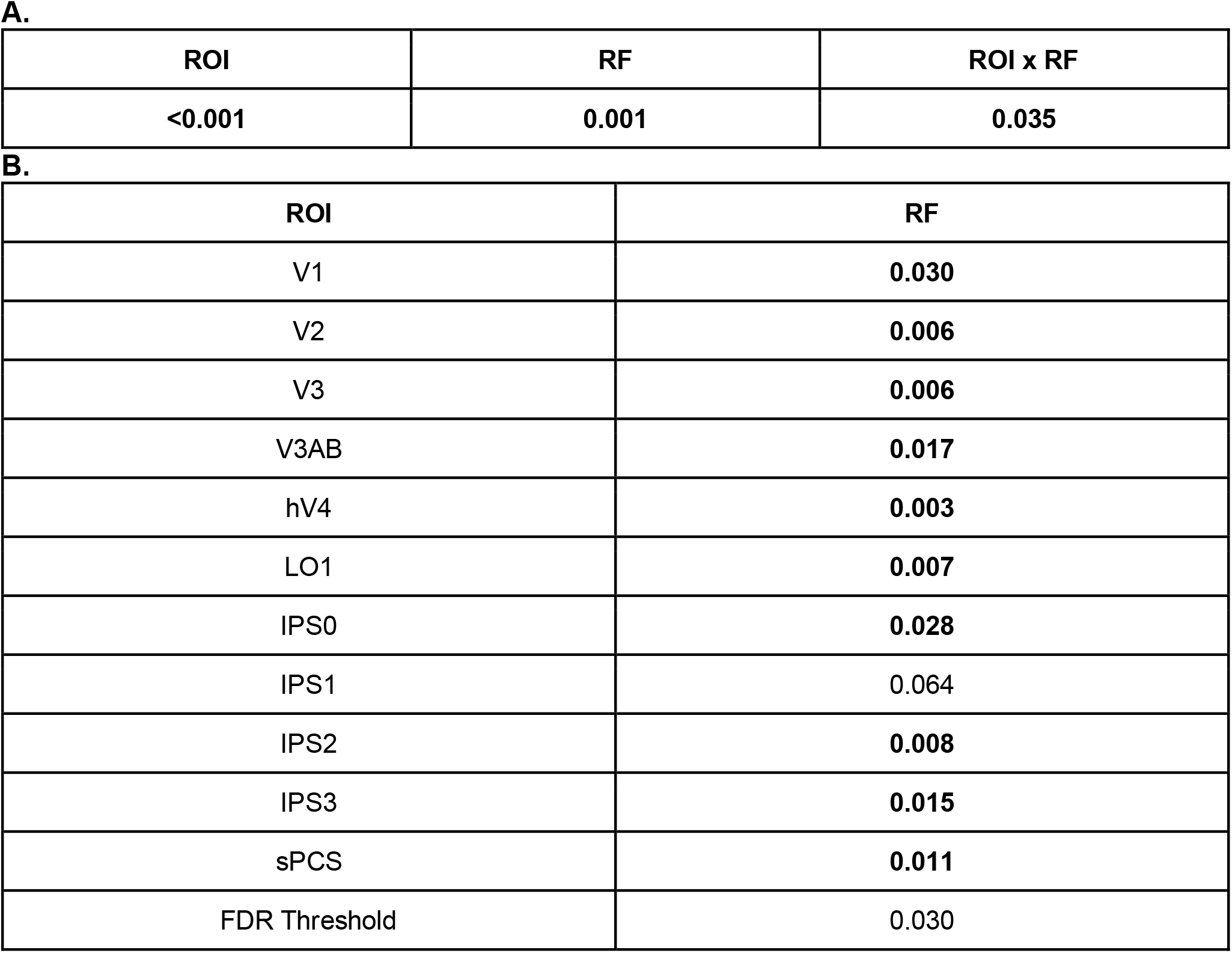
Non-parametric *p-*values characterizing differences in mean delay period amplitude and RF conditions in individual ROIs, Figure S2. **A**. To test for differences in amplitude across ROIs and reliable differences in RF conditions across dorsal ROIs, a 2 -way shuffled ANOVA was performed on mean delay-period amplitude from distractor-absent trial with ROI and RF conditions as main effects. **B**. A follow-up 1-way shuffled ANOVA was performed within each ROI with RF condition (in vs. out) as factors. Significant tests are marked in bold, trends (*p*<0.05, uncorrected) are marked in italics.

**Table S2.**
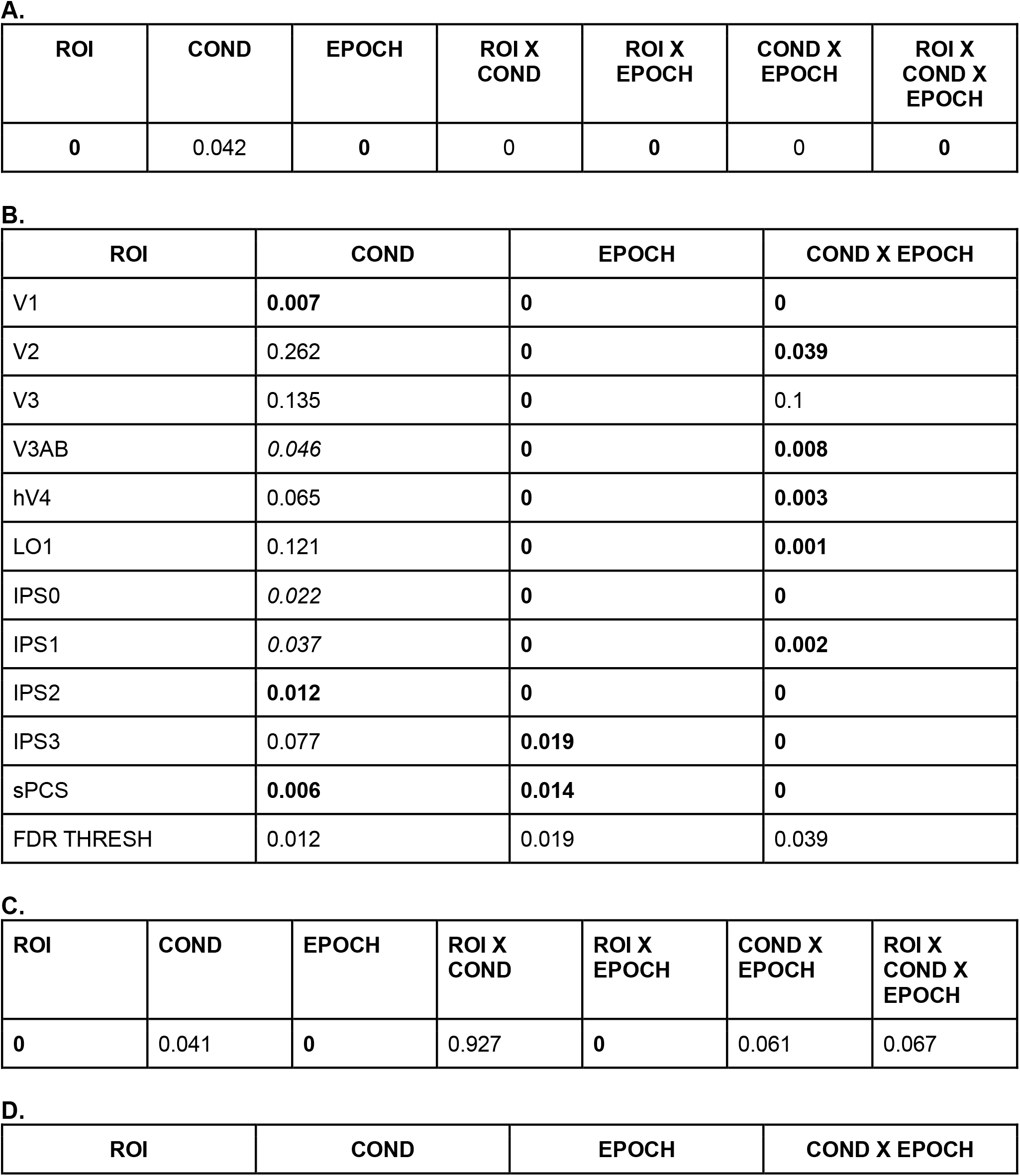
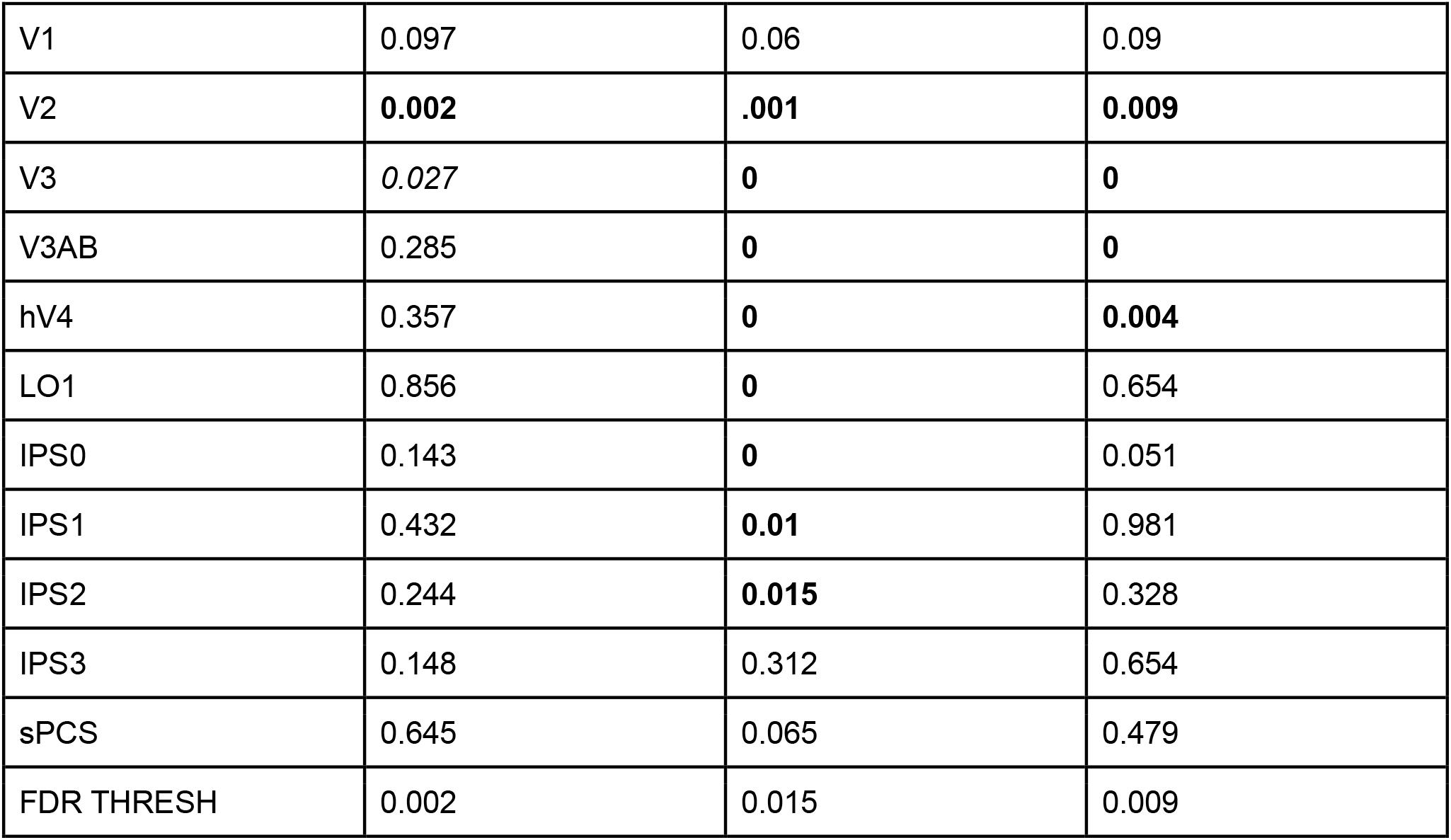
Non-parametric *p*-values characterizing BOLD Z-score averages in individual ROIs, Figure 2. **A**. 3-way ANOVA was performed on RF-in data with ROI, condition, & epoch as main effects. **B**. 2-way permuted ANOVA tests on RF-in data per ROI with condition (distractor absent vs. present) and epoch as main effects. **C**. 3-way ANOVA was performed in RF-in-out data with ROI, condition, & epoch as main effects. **D**. 2-way permuted ANOVA tests on RF-in-out data per ROI with condition (distractor absent vs. present) and epoch as main effects. Significant tests are marked in bold, trends (*p*<0.05, uncorrected) are marked in italics.

**Table S3.** Non-parametric *p*-values characterizing effect of distraction on WM representation fidelity (individual ROIs), Figure S4. p-values for 3-way, 2-way, and *t*-tests per each ROI from Figure S4. For each test, test statistics (F- or T-scores) computed using intact data labels were compared against those computed using data labels shuffled within each participant 1000x.

| Epoch | Cond | ROI | Epoch x Cond | Epoch x ROI | Cond x ROI | Epoch x Cond x ROI |
| --- | --- | --- | --- | --- | --- | --- |
| **< 0.001** | **0.002** | **< 0.001** | **0.001** | **< 0.001** | 0.479 | **0.028** |

|  | Epoch | Condition | Epoch x Condition |
| --- | --- | --- | --- |
| V1 | 0.689 | **0.004** | 0.202 |
| V2 | *0.05* | **0.007** | **0.006** |
| V3 | **0.001** | **0.005** | **< 0.001** |
| V3AB | **< 0.001** | **0.005** | **< 0.001** |
| hV4 | **0.001** | **0.008** | *0.048* |
| LO1 | **0.003** | **0.017** | 0.722 |
| IPS0 | **< 0.001** | **0.005** | **0.004** |
| IPS1 | **0.023** | **0.017** | *0.031* |
| IPS2 | 0.109 | **0.001** | 0.358 |
| IPS3 | 0.388 | **0.004** | 0.272 |
| sPCS | 0.361 | **0.033** | 0.089 |
| FDR Threshold | 0.023 | **0.033** | 0.006 |

|  | Epoch 1 (PRE) | Epoch 2 (DIST) | Epoch 3 (POST) |
| --- | --- | --- | --- |
| V1 | **0.0260** | **0.0120** | **0.0060** |
| V2 | 0.1160 | **0.0020** | **0.0080** |
| V3 | 0.1600 | **< 0.001** | *0.0360* |
| V3AB | 0.2340 | **0.0020** | *0.0360* |
| hV4 | **0.0160** | **0.0040** | *0.0500* |
| LO1 | **0.0040** | **0.0180** | *0.0420* |
| IPS0 | 0.1940 | **< 0.001** | *0.0460* |
| IPS1 | 0.0660 | **0.0020** | **0.0140** |
| IPS2 | **0.0020** | **< 0.001** | **0.0300** |
| IPS3 | **0.0020** | **< 0.001** | *0.0500* |
| sPCS | 0.0820 | **0.0020** | 0.1420 |
| FDR THRESH | 0.03 | **0.03** | 0.03 |

**Table S4.** Non-parametric *p*-values characterizing effect of distraction on WM representation fidelity, Figure 5B. *p*-values for 3-way, 2-way, and *t*-tests per each ROI from Figure 5B. For each test, test statistics (F- or T-scores) computed using intact data labels were compared against those computed using data labels shuffled within each participant 1000x.

| Epoch | Condition | ROI | Epoch x Condition | Epoch x ROI | Cond x ROI | Epoch x Condition x ROI |
| --- | --- | --- | --- | --- | --- | --- |
| **< 0.001** | **0.001** | **< 0.001** | **0.001** | **< 0.001** | 0.551 | **0.041** |

|  | Epoch | Condition | Epoch x Condition |
| --- | --- | --- | --- |
| V1-V3 | **0.013** | **0.011** | **0.002** |
| V3AB | **0** | **0.004** | **0.001** |
| hV4 | **0.001** | **0.008** | *0.044* |
| LO1 | **0.002** | **0.011** | 0.747 |
| IPS0/1 | **0.001** | **0.002** | **0.004** |
| IPS2/3 | 0.187 | **0.002** | 0.181 |
| sPCS | 0.334 | **0.036** | 0.087 |
| FDR THRESH | 0.013 | 0.036 | 0.004 |

|  | Epoch 1 (PRE) | Epoch 2 (DIST) | Epoch 3 (POST) |
| --- | --- | --- | --- |
| V1-V3 | 0.076 | **0.004** | **0.012** |
| V3AB | 0.218 | **0.002** | **0.022** |
| hV4 | **0.026** | **0.004** | 0.056 |
| LO1 | **0.006** | **0.01** | *0.042* |
| IPS0/1 | 0.096 | **0.002** | **0.006** |
| IPS2/3 | **< 0.001** | **< 0.001** | **0.03** |
| sPCS | 0.124 | **< 0.001** | 0.112 |
| FDR THRESH | 0.03 | 0.03 | 0.03 |

**Table S5.** Non-parametric *p*-values quantifying impact of model estimation procedures (individual ROIs), Figure S5. Statistics for Figure S5C (comparison of WM fidelity on distractor-present trials across model estimation procedures). A. A 3-way ANOVA was performed with ROI, model, & epoch as factors For each ROI, we performed a 2-way repeated measures ANOVA against a shuffled null (within each participant, shuffle datapoint labels 1000x). For each test (main effect of model, main effect of epoch, interaction), we compute an FDR threshold. Significant tests are marked in bold, trends (*p*<0.05, uncorrected) are marked in italics.

**A. All ROIs 3-way ANOVA:**
| ROI | Model | Epoch | ROI × Model | ROI × Epoch | Model × Epoch | ROI × Model × Epoch |
| --- | --- | --- | --- | --- | --- | --- |
| < 0.001 | 0.005 | <0.001 | 0.326 | <0.001 | 0.055 | 0.001 |

**B. 2-way repeated measures ANOVA:**
| Figure S5C | Model | Epoch | Interaction |
| --- | --- | --- | --- |
| V1 | 0.147 | 0.259 | 0.934 |
| V2 | 0.031 | **0.003** | 0.238 |
| V3 | *0.023* | <0.001 | *0.015* |
| V3AB | 0.054 | <0.001 | *0.021* |
| hV4 | 0.02 | <0.001 | 0.179 |
| LO1 | *0.018* | <0.001 | 0.075 |
| IPS0 | **0.009** | **0.001** | 0.121 |
| IPS1 | <0.001 | **0.010** | 0.866 |
| IPS2 | 0.239 | **0.036** | 0.288 |
| IPS3 | 0.226 | 0.146 | 0.335 |
| sPCS | 0.065 | 0.084 | 0.176 |
| FDR THRESHOLD | 0.009 | 0.036 | <0.001 |

**Table S6.** Non-parametric *p*-values quantifying impact of model estimation procedures, Figure 6. Statistics for Figure 6D (comparison of WM fidelity on distractor-present trials across model estimation procedures). For each ROI, we performed a 2-way repeated measures ANOVA against a shuffled null (within each participant, shuffle datapoint labels 1000x). For each test (main effect of model, main effect of epoch, interaction), we compute an FDR threshold. Significant tests are marked in bold, trends (*p*< 0.05, uncorrected) are marked in italics. Related to Figure 6D.

| ROI | Model | Epoch | ROI x Model | ROI x Epoch | Model x Epoch | ROI x Model x Epoch |
| --- | --- | --- | --- | --- | --- | --- |
| **< 0.001** | **0.010** | **<0.001** | 0.641 | **<0.001** | **0.029** | **0.018** |

| Figure 6D | Model | Epoch | Model x Epoch |
| --- | --- | --- | --- |
| V1-V3 | *0.037* | **0.001** | 0.127 |
| V3AB | *0.047* | **<0.001** | *0.017* |
| hV4 | *0.023* | **<0.001** | 0.145 |
| LO1 | *0.015* | **<0.001** | 0.079 |
| IPS0/IPS1 | **0.004** | **<0.001** | 0.097 |
| IPS2/IPS3 | 0.248 | *0.047* | 0.196 |
| sPCS | *0.048* | 0.086 | 0.212 |
| FDR thresh | 0.004 | 0.001 | **<0.001** |

**Table S7.**
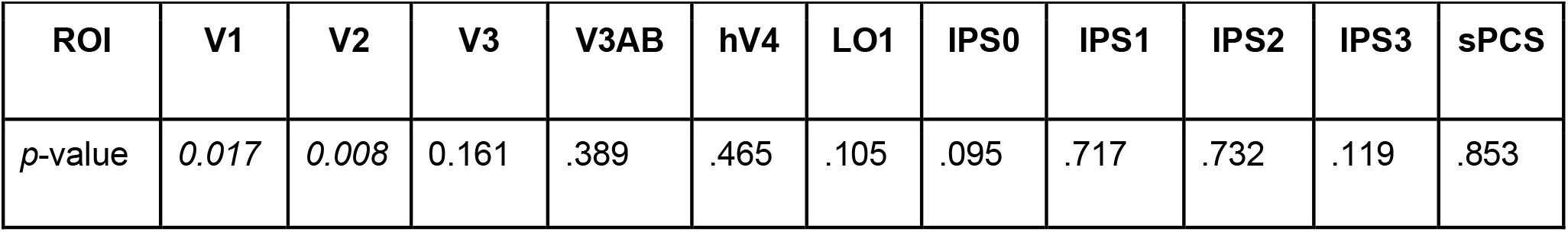
P-values characterizing neural-behavioral correlations (individual ROIs), Figure S6B. Shuffled correlations were performed on an individual basis and subjected to t-test on Fisher r-to-Z transformed correlation values, 1000x. No ROIs pass FDR correction. Italics indicates trends, defined as p < 0.05, uncorrected.

**Table S8.** P-values characterizing neural-behavioral correlations, Figure 7C. Shuffled correlations were performed on an individual basis and subjected to *t*-test on Fisher r-to-Z transformed correlation values, 1000x. FDR threshold is *p*=0.005. Bold indicates significant ROIs, corrected for multiple comparisons.

| ROI | V1-V3 | V3AB | hV4 | LO1 | IPS0/1 | IPS2/3 | sPCS |
| --- | --- | --- | --- | --- | --- | --- | --- |
| *p*-value | **0.005** | 0.4104 | 0.4710 | 0.1040 | 0.4310 | 0.3130 | 0.8670 |

## STAR METHODS

### KEY RESOURCES TABLE

| REAGENT or RESOURCE | SOURCE | IDENTIFIER |
| --- | --- | --- |
| Software and Algorithms |  |  |
| MATLAB 2018a | MathWorks | https://www.mathworks.com |
| PsychToolBox | psychtoolbox.org | http://psychtoolbox.org/ |

## RESOURCE AVAILABILITY

### Lead Contact

Further information and requests for resources and reagents should be directed to and will be fulfilled by the Lead Contact, Dr. Clayton Curtis

### Materials Availability

This study neither used any reagents nor generated any new materials

### Data and Code Availability

All data supporting conclusions in this report [will be made] freely available on the Open Science Framework (URL), and code used to present stimuli, analyze data and produce figures will be made freely available on GitHub (github.com/clayspacelab). For participant privacy reasons, for fMRI data we only publicly provide extracted activation timecourses from each ROI for each participant, as well as each voxel’s best-fit pRF parameters and extracted timecourses. Stimulus presentation code for retinotopic mapping using Psychtoolbox is available on GitHub (github.com/clayspacelab/vRF_stim). Additionally, code used to preprocess raw data can be inspected on GitHub. We preprocessed fMRI data using shell scripts and custom MATLAB functions available on GitHub (github.com/tommysprague/preproc_shFiles and github.com/tommysprague/preproc_mFiles), and we fit voxel receptive fields using a GPU-accelerated branch of mrVista, available on GitHub (github.com/tommysprague/vistasoft_ts), which implements GPU-accelerated gridfits (github.com/tommysprague/gridfitgpu).

## EXPERIMENTAL MODEL AND SUBJECT DETAILS

### Participants

Seven neurologically healthy volunteers (3 female; 25-50 years old) with normal or corrected-to-normal vision participated in this study after giving informed consent, using procedures approved by New York University IRB (protocol IRB-FY2017-1024). Each participant completed 1-2 sessions of retinotopic mapping and anatomical scans (∼2hrs), between 2-3 independent mapping sessions for model training (∼1.5hrs each), and two experimental sessions (∼1.5hrs each). Sample size was chosen based on similar sample sizes recruited for similar studies employing deep, multi-session imaging per participant (Rademaker et al., 2019, n=6; Sprague et al., 2016, n=6).

## METHOD DETAILS

### Stimulus

Stimuli were generated via PsychToolBox in Matlab 2018b on a PC and presented via a contrast-linearized ViewPixx ProPixx projector, with resolution 1280×1024 and refresh rate 120Hz. Participants viewed the stimuli through a mirror attached to the head coil at a viewing distance of 63cm. The projected image spanned 36.3cm height, resulting in a maximum field of view of 32.14° visual angle.

### Experimental task

Participants performed a modified version of the memory guided saccade task (Hikosaka and Wurtz, 1983) (Figure 1A-B). All stimuli were presented within a 15° radius circular grey aperture on a black background. Throughout the whole experiment, a 0.075° radius light gray fixation point was presented in the center of the aperture. We tested two conditions, randomly interleaved: distractor-absent trials required participants to precisely remember a target location over an extended delay interval, and distractor-present trials required participants to additionally perform a visual motion discrimination task based on a peripheral visual stimulus. Each trial began with a pre-cue (1000ms; 0.55° circle centered at the fixation point) reliably informing participants whether a distractor would (cyan) or would not (magenta) be presented during the delay period. This controlled for possible impacts of distractor predictability on cognitive control of WM representations (Bettencourt and Xu, 2016; Rademaker et al., 2019). Next, a light gray target dot (0.65° diameter) was presented at a randomly-chosen position along an invisible ring 12° around fixation for 500ms. Participants precisely remembered the target location over the subsequent 12s delay interval. On distractor-absent trials, no further stimuli appeared during the delay period. On distractor-present trials, 4,500ms into the delay period a distracting stimulus appeared at one of seven positions relative to the WM target position around the invisible 12° eccentricity ring for 1,000ms. On each distractor-present trial, distractor positions were randomly chosen (counterbalanced within a run), and were further jittered by ±12° polar angle. The distractor stimulus was a random dot kinematogram (RDK) containing equal numbers of black and white dots (100% contrast; 17 dots/deg^2^, 0.075° dot radius, 1° radius of dot patch; 0.1s dot lifetime), with a subset of dots rotating either clockwise or counterclockwise about the center of the dot patch (∼100° polar angle/s). Participants responded within 2.5s of the distracting stimulus onset whether the coherently moving dots rotated clockwise (right button) or counterclockwise (left button). The coherence of the dot patch was fixed within a run, and was adjusted between runs based on behavioral accuracy to achieve ∼75% performance (mean = 73%, SEM = 6.74%). Non-coherent dots each moved in a random direction. The remainder of the delay (6,500ms) was identical to the distractor-absent trials.

On all trials, at the end of the full 12s delay period, the fixation point was replaced with a filled light gray circle, cueing participants to make a saccade towards the remembered position. After 800ms, the target dot was re-presented for 1,000ms, which participants were instructed to fixate. Each trial ended with a 7 to 13s ITI (in 750ms steps), randomly chosen. In addition, on distractor-present trials, participants received feedback on their discrimination performance at the beginning of the ITI (red or green circle at fixation for correct or incorrect response, respectively). Across two sessions, participants performed between 25-36 runs (12-18 runs per session, 279s per run), where each run consisted of seven distractor-present and three distractor-absent trials, randomly ordered. This resulted in between 250 and 360 total experimental trials per participant.

### Independent model estimation task

We acquired a separate, independent dataset used only to estimate the inverted encoding model for spatial position. Participants performed a simple memory-guided saccade task over a 12s delay using a nearly identical stimulus display, but with no distractor stimulus. Within each 16-trial run, target locations were chosen from a discrete set of positions spaced evenly around an invisible ring (12° eccentricity), with the set of positions staggered every other run, for 32 unique positions overall. No pre-cue was presented at the beginning of each trial. Participants performed between 20 and 31 runs, across 2-3 separate scanning sessions, where each run was composed of 16 trials and lasted 396s.

### Retinotopic mapping task

To identify regions of interest (ROIs) for all reported analyses, we acquired retinotopic mapping data using an adaptation of a previously-reported task and procedure (Mackey et al., 2017) and computed polar angle and eccentricity maps using the population receptive field (pRF) method (Dumoulin and Wandell, 2008). During each retinotopy run, subjects completed a difficult discrimination task within bars that swept across 26.4° of the visual field in twelve 2.6s steps. Bar widths and sweep directions were pseudo-randomly chosen from three different widths (2.5°, 5.0°, and 7.5°) and four directions (left-to-right, right-to-left, bottom-to-top, top-to-bottom), respectively. Each bar was split into three equally-sized rectangular patches along its long axis. Each of the three patches contained an RDK moving in one of the two possible directions perpendicular to the bar’s swipe direction (parallel to the long axis). During each step of the swipe, the RDK direction in one of the peripheral patches matched the RDK direction in the central patch. Subjects were instructed to report, via a button press, which peripheral patch matched the central patch’s RDK direction, at each step of the swipe. After each report, participants received accuracy feedback (red or green fixation point), and a three-down/one-up staircase was implemented to maintain task difficulty ∼80%. The coherence of the RDK in two peripheral patches was always 50% while a variable RDK coherence was used in the central patch to adjust the difficulty of the task. Stimulus presentation code is available at github.com/clayspacelab/vRF_stim

## QUANTIFICATION AND STATISTICAL ANALYSIS

### fMRI acquisition

All functional MRI images were acquired at the NYU Center for Brain Imaging 3T Siemens Prisma Scanner. fMRI scans for experimental, model estimation, and retinotopic mapping were acquired using the CMRR MultiBand Accelerated EPI Pulse Sequences (Release R015a) (Moeller et al., 2010); (Feinberg et al., 2010); (Xu et al., 2013). All functional and anatomical images were acquired with the Siemens 64 channel head/neck coil.

### Experimental & model estimation scans

For the experimental and model estimation scans, BOLD contrast images were acquired using a Multiband (MB) 2D GE-EPI with MB factor of 4, 44 2.5mm interleaved slices with no gap, isotropic voxel size 2.5mm and TE/TR: 30/750ms. We measured field inhomogeneities by acquiring spin echo images with normal and reversed phase encoding (3 volumes each), using a 2D SE-EPI with readout matching that of the GE-EPI and same number of slices, no slice acceleration, TE/TR: 45.6/3537ms.

### Retinotopic mapping scans

For the retinotopic mapping scans, BOLD contrast images were acquired using a Multiband (MB) 2D GE-EPI with MB factor of 4, 56 2mm interleaved slices with no gap, isotropic voxel size 2mm and TE/TR: 42/1300ms. Distortion mapping scans were acquired with normal and reversed phase encoding, using a 2D SE-EPI with readout matching that of the GE-EPI and same number of slices, no slice acceleration, TE/TR: 71.8/6690ms.

### Anatomical and bias images

T1- and T2-weighted images were acquired using the Siemens product MPRAGE and Turbo Spin-Echo sequences (both 3D) with 0.8 mm isotropic voxels, 256 × 240 mm slice FOV, and TE/TR of 2.24/2400 ms (T1w) and 564/3200 ms (T2w). We collected 192 and 224 slices for the T1w and T2w images, respectively. We acquired between two and five T1 images, which were aligned and averaged to improve signal-to-noise ratio. In addition, to correct functional images for inhomogeneities in the receive coil sensitivity and improve the motion correction and coregistration process, we collected two fast 3D GRE sagittal images (resolution: 2mm isotropic, FoV: 256 × 256 × 176 mm; TE/TR: 1.03/250 ms), one with the body coil and the other with the 64 ch head/neck coil.

### fMRI preprocessing

We used all intensity-normalized high-resolution anatomical scans (for each participant, 2-5 T1 images and 1 T2 image) as input to the ‘hi-res’ mode of Freesurfer’s recon-all script (version 6.0) to identify pial and white matter surfaces. We edited these surfaces by hand using Freeview as necessary and converted surfaces to SUMA format. The processed anatomical image for each participant acted as the alignment target for all functional datasets. Our aim for functional preprocessing was to put functional data from each run into the same functional space at the same voxel size acquired during the task sessions (2.5mm isovoxel), account for run- and session-specific distortions, incur minimal volume-wise smoothing by minimizing spatial transformations, and apply a marginal amount of smoothing along the direction orthogonal to the cortical surface. This allowed us to optimize SNR and minimize smoothing, ensuring ROI data remains as near as possible to its original dimensionality. Moreover, because the distortion field can depend on the exact position of the head within the main field, we divided functional sessions into 3-5 ‘mini-sessions’ consisting of 1-4 task runs split by a pair of spin-echo images acquired in opposite phase encoding directions, used for anatomical registration and computing distortion fields for distortion correction. We applied all preprocessing steps described below to each mini-session independently, and inspected motion correction, coregistration and distortion correction to ensure the procedures worked as intended. Preprocessing was performed using a combination of scripts generated with AFNI’s afni_proc.py and custom scripts implementing AFNI functions (version 17.3.09, pre-compiled Ubuntu 16 64 bit distribution). We performed all analyses on a LINUX workstation running Ubuntu v16.04.1 using 8 cores for most OpenMP accelerated functions.

First, we corrected functional images for intensity inhomogeneity induced by the high-density receive coil by dividing all images by a smoothed bias field (15mm FWHM), computed as the ratio of signal in the receive field image acquired using the head coil to that acquired using the in-bore ‘body’ coil. To improve coregistration of functional data to the target T1 anatomical image, we used distortion-corrected and averaged spin-echo images (which were used to compute distortion fields restricted to the phase-encode direction) to compute transformation matrices between functional and anatomical images. Then, we used the rigorous distortion-correction procedure implemented in afni_proc.py to undistort and motion-correct functional images. Briefly, this procedure involved first distortion-correcting all images in each run using the distortion field computed from the spin-echo image pair, then computing motion-correction parameters (6-parameter affine transform) using these unwarped images. Next, we used the distortion field, motion correction transform for each volume, and the functional-to-anatomical coregistration simultaneously to render functional data from native acquisition space into unwarped, motion corrected, and coregistered anatomical space for each participant at the same voxel size as data acquisition in a single transformation and resampling step. For retinotopic mapping data, this was a 2mm isovoxel grid; and for task data, this was 2.5mm isovoxel grid. For both task and retinotopy data, we projected this volume-space data onto the reconstructed cortical surface. For retinotopy data, we made a smoothed version of the data by smoothing on the cortical surface (5mm FWHM). We then projected surface data (for task data, only the ‘raw’ data; for retinotopy data, the raw and smoothed data) back into volume space for all analyses. For unsmoothed data, this results in a small amount of smoothing for each voxel along a vector orthogonal to the surface in volume space.

To compute pRF properties (see below) in the same voxel grid as task data, we projected retinotopy time series data onto the surface from its native space (2mm iso), then from the surface to volume space at the task voxel resolution (2.5mm iso). This ensured that variance explained estimates faithfully reflect goodness of fit and are not impacted by smoothing incurred from transforming fit parameter values between different voxel grids. We linearly detrended activation values from each voxel from each run and converted signal to percent signal change by dividing by the mean over the entire run. For multivariate analyses on task data, we subsequently Z-scored each voxel across all volumes for each run independently.

### Retinotopic mapping and ROI definition

We averaged time series from each voxel across all retinotopy runs (9-12 per participant) in volume space and fit a pRF model for each voxel using a GPU-accelerated extension of vistasoft (github.com/clayspacelab/vistasoft). We fit a compressive spatial summation isotropic Gaussian model (Kay et al., 2013); (Mackey et al., 2017) as implemented in mrVista (see (Mackey et al., 2017) for detailed description of the model). We created a high-resolution stimulus mask (270 × 270 pixels) to ensure similar predicted response within each bar size across all visual field positions (to mitigate the effects of aliasing with a lower-resolution stimulus mask grid), and began with an initial high-density grid search, followed by subsequent nonlinear optimization. Note that, in these analyses, because we conduct a grid search on all voxels independently, there is no smoothing of parameter estimates applied after this step before nonlinear optimization. For all analyses described below, we used best-fit pRF parameters from this nonlinear optimization step.

After estimating pRF parameters for every voxel in the brain, ROIs were delineated by projecting pRF best-fit polar angle and eccentricity parameters with variance explained ≥10% onto each participant’s inflated surfaces via AFNI and SUMA. ROIs were drawn on the surface based on established criteria for polar angle reversals and foveal representations (Mackey et al., 2017); (Wandell et al., 2007); (Winawer and Witthoft, 2015); (Amano et al., 2009); (Swisher et al., 2007). Finally, ROIs were projected back into volume space to select voxels for analysis. In this report we consider data from V1, V2, V3, V3AB, hV4, LO1, IPS0, IPS1, IPS2, IPS3, and sPCS, which are all retinotopic regions described in previous reports examining the impact of visual distractors on WM representations (Bettencourt and Xu, 2016; Lorenc et al., 2018; Rademaker et al., 2019). Only voxels with ≥10% variance explained in pRF model fits were included in subsequent fMRI analyses. Additionally, we group ROIs V1-V3, IPS0-1, and IPS2-3 by concatenating voxels before multivariate analyses for results presented in Figs 3-7 because they belong to clustered defined by overlapping foveal representations (Wandell et al., 2007), and for consistency with a prior report (Lorenc et al., 2018); see Supplementary Material for analyses reported for each ROI individually).

### Oculomotor Processing

Monocular eye-tracking data were recorded using an MR-compatible Eyelink 1000 infrared eye tracker (SR Research). Eye position (X,Y) and pupil size were recorded at 500 Hz. Prior to the beginning of the experiment, eye position was calibrated with a 13-, or 9-point calibration. Eye data were transformed from raw pixel screen coordinates into degrees of visual angle utilizing the freely available iEye toolbox (github.com/clayspacelab/iEye_ts). First, the data was rescaled given known screen resolution and viewing distance. Next, the data was inspected for extreme values and blinks, which were removed. It was then smoothed with a Gaussian kernel (5ms SD). Saccades were identified by identifying periods with velocity in excess of 30°/s for at least 7.5ms, and resulting in at least a 0.25° amplitude gaze shift. Then, the data for an entire trial was drift corrected by taking the mean over known epochs when the participant is fixating and subtracting that value from the entire trial. Finally, the data is recalibrated to the target position on a run-wise basis by fitting a 3^rd^ order polynomial to X and Y eye positions independently to best approximate the true target location on each trial. On a given trial, if the initial saccade did not meet the following additional criteria, that trial was also excluded from behavioral analysis, and from correlations with neural data (Figure 7): less than a total duration of 150ms, at least 5° in amplitude, and within at least 5° error from the target location. Trials could additionally be removed from analysis if the participant exhibited a fixation break of at least 2.5° during the WM delay or did not make a saccade within the specific response epoch.

### Oculomotor Analysis

Once saccades were preprocessed, we used the last endpoint of the memory-guided saccade before the reappearance of the target, after any corrective saccades, as a measure of the participant’s behavioral WM report in our analyses. For all analyses, we realigned all visual field coordinates of both memory-guided saccade and distractor locations with reference to the saccade target location. We subtracted the polar angle of the memory target from the polar angle of the memory-guided saccade while keeping the radius of each location untouched (to preserve the eccentricity of the reported position). For our precision analysis, we calculated each participant’s saccade standard deviation (SD) by computing the across-trial standard deviations of saccadic polar angle. We calculated reaction times by taking the time from the onset of the response cue to the onset of the initial saccade.

### fMRI: univariate analysis

To assess the mean response across spatially-selective voxels subtending relevant locations (target/distractor) in all ROIs, we computed an event-related average of measured BOLD response for each condition separately. After extracting Z-scored BOLD signal (see Preprocessing) from each voxel, we sorted voxels on each trial according to their best-fit pRF parameters and the known location(s) of the target and/or distractor. RF-in responses (corresponding to voxels tuned nearby the relevant location) were determined by selecting voxels with ≥ 10% variance explained, eccentricity between 2 and 15 degrees, and polar angle difference between the WM target (or distractor) and pRF center of each voxel ≤ 15 degrees. RF-out responses were determined by selecting voxels with ≥ 10% variance explained, eccentricity between 2 and 15 degrees, and a polar angle difference between the WM target (or distractor) and pRF center of each voxel ≥ 165 degrees. We averaged responses across such selected voxels within each ROI, then across all trials within a condition (Figure 2, all ROIs in Figure S2).

Additionally, we removed the baseline response measured between -2.25 and 0 s relative to delay onset.

### fMRI: multivariate inverted encoding model

To reconstruct the representation of visual polar angle carried by the neural population activity of each ROI at any given time, we implemented an inverted encoding model (IEM) (Brouwer and Heeger, 2009; Sprague et al., 2018b) which maps between a modeled encoding space and the measured response across a set of fMRI voxels using a simplified set of information channels (or basis functions) each with a preferred stimulus value. The model assumes that the activation of each voxel in response to a stimulus can be described as a simplified encoding model built via a linear combination of all modeled channel responses to that stimulus. Based on this assumption, the activation of these modeled channels most likely to give rise to an observed activation pattern can be estimated using the inverse of the encoding model. Thus, we can map one space to the other by estimating the strength of the links (i.e. regression weights) between all voxels in a given region and all of the modeled information channels. Note that this analysis framework does not, and cannot, infer ‘tuning’ properties of neurons within the regions analyzed (Gardner and Liu, 2019). Instead, it recovers region-level representations of a feature space (here, spatial position parameterized by polar angle). See (Sprague et al., 2018b) and (Sprague et al., 2019) for a detailed discussion.

To build an IEM for each ROI, we first obtained an independent dataset which we used to calculate the regression weights that describe the encoding model for each voxel (one weight for each channel for each voxel). Model estimation data was always the average delay-period activation between 5.25-12s following delay onset (average over 9 TRs, each 750ms). While these regression weights enable us to map the stimulus space to the voxel space, we used inverted weights to estimate each channel’s response and used them to reconstruct the stimulus space given the population activity pattern at each timepoint. Through this method, we were able to reconstruct the spatial representation of the memory target and the distractor location from the neural population activity during WM. The linear mapping between the voxel space and the stimulus space is defined as:

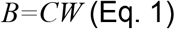

Where, *B* is a matrix (*n* trials × *m* voxels) consisting of the activity of all voxels in a given ROI across trials, *C* is a matrix (*n* trials × *k* channels) containing the response of all channels across the same trials, and *W* is a matrix (*k* channels × *m* voxels) of regression weights describing how much each modeled channel contributes to the BOLD signal measured in each voxel. As the stimuli always appeared along a fixed annulus of 12° in this experiment, we modeled the information channels as 1-D smooth tuned filters centered at 8 uniformly distributed polar angles (ψ) around the 360° polar space (Figure 2):

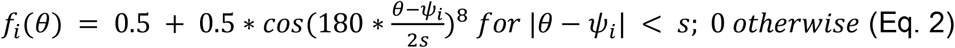

We used a size constant (s) of 180°. To calculate each channel output from the population activity in response to a given stimulus, we first need to calculate the regression weights (*W*) in (Eq. 1). We can do this using measured activation patterns (*B*_*trn*_) and corresponding predicted channel responses (*C*_*trn*_) for model estimation data:

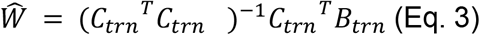

Where *B*_*trn*_ is a matrix of BOLD activity in response to our training stimuli (trials × voxels) and *C*_*trn*_ is a matrix containing the corresponding predicted channel outputs calculated from (Eq. 2), given the training stimulus locations (polar angle). For the IEM, we used these regression weights calculated from an independently collected training set to estimate channel outputs from the measured BOLD signal at any given epoch in our WM task. We estimated these channels outputs as:

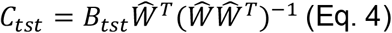

Where *B*_*tst*_ contains the measured BOLD signal on each time point of experimental trials, Ŵ contains the regression weights (estimated using an independent dataset, Eq. 3), and *C*_*tst*_ is the estimated channel responses.

Finally, to reconstruct the stimulus space, given a set of computed channel responses (per timepoint, epoch, or trial), we calculate the sum of all channel sensitivity profiles, each weighted by its corresponding estimated channel response. For visualization, we align each trial based on the known target location (Figures 2C-D, 3A-B) or the known distractor location (Figures 2E-F, 3C) by circularly shifting the reconstruction such that the aligned position is denoted as 0°. Note that for distractor-present trials, the reconstruction necessarily contains both a representation of the distractor and a representation of the target location. But, because these were randomized with respect to one another, aligning to one results in the other representation ‘averaging out’ (Figure 2).

### Model training

For Figures 2-5, we estimated the IEM using delay-period activation (5.25-12 s) measured from an independent mapping dataset in which participants performed a single-item MGS task (see above). Additionally in Figure 5, we tested whether a model estimated using distractor-present trials contains WM information in a neural format different from that used in the independent mapping dataset. We performed a leave-one-run-out cross-validation procedure whereby all distractor-present trials from all runs but one (concatenated across sessions) are used to estimate the IEM, and this IEM is used to reconstruct WM representations from the held-out run. We repeat this procedure over all runs, until each distractor-present trial has served as a ‘test’ trial.

Moreover, to establish whether the format of WM representations transforms over the delay period following disruption by a distractor (e.g., (Parthasarathy et al., 2017; Spaak et al., 2017), we performed this leave-one-run-out analysis for each pair of training/testing timepoints during the trial (Figure 5A), and for each trial epoch (pre-distractor, distractor, post-distractor; Figure 5B-C).

### Fidelity

To quantify the presence of information about a remembered or viewed stimulus location in IEM-based reconstructions, we computed a model-free index of ‘fidelity’ (Rademaker et al., 2019; Rahmati et al., 2020; Sprague et al., 2016). Conceptually, this metric measures whether the reconstruction, on average, ‘points’ in the correct direction, along with how ‘strong’ the representation is. Accordingly, we compute the vector mean of the reconstruction in polar coordinates. When reconstructions are rotated and aligned to 0°, projecting this vector mean on the horizontal axis captures energy in the reconstruction consistent with the aligned location. Each reconstruction r(*θ*), where *θ* is the polar angle of each point and r(*θ*), is the reconstruction activation) when plotted as a polar plot, was projected along the x-axis (reconstructions were rotated such that the target was presented at 0°):

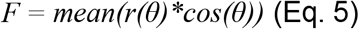

If *F* is reliably greater than zero, this quantitatively demonstrates that the net activation over the entire reconstruction carries information above chance about the target position. This measure is independent of baseline activation level in the reconstruction, as the mean of *r(θ)* is removed by averaging over the full circle. We computed fidelity for each timepoint of each trial in the experiment after aligning reconstructions to the target position, and, on distractor-present trials, the distractor position. Because the non-aligned stimulus type (target or distractor) was located at a random and symmetric position with respect to the aligned stimulus type (Figure 1A), we could independently assay information about both stimulus types on the same set of trials (see also (Rademaker et al., 2019)).

### Quantifying decoded position

We also used single-trial reconstructions on trials with distractor locations nearby the WM target location to decode the position (polar angle) represented by the population activation pattern. We defined the decoded position (*WM*_*est*_) as the circular mean of the reconstruction, which we computed by summing over unit vectors pointing in each direction weighted by the reconstruction activation in that direction, then taking the inverse tangent of the resulting vector:

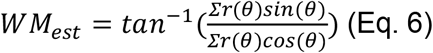

### Statistical Procedures

To test whether behavioral performance was impacted by distractor presence or absence, we performed one-way repeated measures ANOVAs on memory error and saccadic reaction times. Memory error was defined as the standard deviation of target-aligned saccadic polar angle (degrees). Reaction time was defined as time at which an initial ballistic saccade was made in the response period, as measured from the onset of the response period (milliseconds).

To demonstrate the presence of information about the target/distractor location in an IEM-based reconstruction, we computed *t*-statistics on each fidelity timepoint (Figure 3D-E) by first creating an empirical null *t*-distribution from shuffling trial labels of the training dataset within each participant 1000x. The sample of fidelity values computed using an intact model at each timepoint was then compared against the T distribution computed using shuffled data. We compared the T-score computed using intact trial labels to this null distribution, and defined the p-value as the proportion of null T values equal to or exceeding the actual value (one-tailed; (Rademaker et al., 2019)). We corrected for multiple comparisons using the false discovery rate across timepoints within each ROI and condition.

To quantify how distraction impacted univariate fMRI responses (Figure 2 & Figure S2), we first conducted a 2-way ANOVA using ROI and RF condition (in vs. out) on average BOLD delay-period (3.75-12s) responses from voxels selected by their proximity to (within 15 degrees polar angle; RFin) or separation from (at least 165 degrees polar angle from) the WM target (Fig. 2, A,B; Fig. S2 A,B) or distractor (Fig 2., C; Fig S2 C). F-scores computed for each main effect were compared against a distribution of shuffled data, within participant 1000x. p-values were determined by the proportion of shuffled F-scores per main effect greater than or equal to the F-score computed using the intact data. We performed follow-up 1-way ANOVAs within each ROI, using RF (in vs. out) as the main effect, and F-scores were computed using the same within-participant shuffling procedure. FDR corrections were performed across ROIs.

To quantify WM representation fidelity (Figure 4) in each ROI during each trial epoch, we first conducted a 3-way repeated measures ANOVA (ROI × distractor condition × delay epoch) on average BOLD responses across voxels within each ROI and trials within each condition. F-scores computed for each main effect, 2-way, and 3-way interaction were compared against a distribution of each comparison computed using data shuffled within each participant over 1000 iterations. P-values reflect the proportion of null F-scores within each comparison greater than or equal to the F-score computed using intact data. The minimum *p*-value achievable with this procedure is 0.001. We conducted follow-up 2-way repeated-measures ANOVAs for each ROI (distractor condition x delay epoch) using the same shuffling procedure within each ROI. P-values were corrected for multiple comparisons across all ROIs within each comparison (separately for each main effect and interaction; FDR).

To establish whether WM representation fidelity differed across distractor conditions within each ROI and epoch (Figure 4A), we used a slightly different shuffling procedure. Computing WM fidelity on distractor-present trials cannot occur on single trials, and instead must aggregate across an equivalent number of trials for each relative distractor position. Each scanning run (10 trials) involved one trial for each relative distractor position, so we were able to compute a mean WM representation fidelity for distractor-absent trials (3) and distractor-present trials (7) on each scanning run (26-36 runs per participant). First, we computed a *t*-statistic from a paired *t*-test for each ROI and epoch to quantify the effect of distractor condition on WM fidelity. Then, we computed a null *t*-distribution after shuffling run-wise average WM fidelity scores within each participant 1000 times. P-values were computed as the proportion of the t-distribution exceeding the t-statistic computed using intact data, and doubled to reflect a two-tailed test. We corrected *p*-values using the false discovery rate across all ROIs and epochs.

To determine whether decoded WM target position just before response onset predicted behavioral response positions on a trial-by-trial basis on distractor-present trials, we extracted decoded WM positions from each ROI (Eq. 6) on each trial with a nearby distractor (within 12° polar angle) and correlated, within each participant, these values with corresponding behavioral responses (Figure 5B-C). We converted Pearson correlation coefficients to Z-scores using the Fisher r-to-Z transform before combining across participants, and computed a t-score for each ROI. To assess significance, we shuffled the relationship between behavioral and neural responses within each participant 1000 times, recomputed correlations for each participant and *t*-scores across participants, then compared the true t-score for each ROI to the corresponding shuffled null distribution. P-values (one-tailed) were corrected across ROIs with FDR. We also tested for a main effect of ROI on average trial-by-trial error correlation across participants using a shuffled 1-way ANOVA.

## Notes

Conflicts of interest: The authors declare no competing financial interests.

### Competing Interest Statement

The authors have declared no competing interest.

